# gRely: Relyability for genome-trained sequence-to-function model predictions

**DOI:** 10.64898/2026.05.23.727431

**Authors:** Abdul Muntakim Rafi, Gokcen Eraslan, Kipper Fletez-Brant

**Affiliations:** AI Biology and Development, Computational Sciences Center of Excellence, Genentech; School of Biomedical Engineering, University of British Columbia

## Abstract

Sequence-to-function (S2F) models predict molecular phenotypes from DNA sequence and are increasingly applied to variant effect prediction (VEP), but their accuracy varies across variants, genes, and tissues. Current practice declares predictions as trustworthy by thresholding on predicted effect magnitude, a crude heuristic that discards low-effect variants where most GWAS signal is expected to act. We present gRely, a meta-model that estimates a score that an individual VEP is correct, from 1,161 features derived from the target variant, gene, tissue, and model. Trained on GTEx fine-mapped eQTLs and Borzoi, gRely reaches 97% sign concordance in its top 20% of scores on held-out variants, genes, and tissues. Critically, it remains discriminative among low-magnitude variants, which effect-size filtering discards; SHAP attribution shows that reliability in the low-effect regime is driven by chromatin context and model cross-replicate agreement rather than effect size. Furthermore, the Borzoi trained meta-model generalizes across other architectures, such as AlphaGenome and Enformer. gRely is the first framework to provide per-prediction confidence scores for S2F models and marks a step toward trustworthy deployment of S2F models in genomic research and clinical applications.

---

Sequence-to-function (S2F) models learn regulatory patterns directly from genomic DNA and predict molecular phenotypes such as gene expression and chromatin state. Beginning with DeepSEA (1), DeepBind (2) and continuing through Basenji (3), BPNet (4), Enformer (5), Sei (6), Borzoi (7), Scooby (8), Decima (9), and AlphaGenome (10) these models have shown that networks trained on large-scale genomic profiles capture important aspects of cis-regulatory logic (11, 12). They are now standard tools for designing cell type-specific promoters, enhancers, and synthetic regulatory sequences (13–16), and are increasingly applied to variant effect prediction (VEP), where the goal is to quantify the functional impact of a genetic variant on regulatory activity (17, 18).

Despite their promise, however, these models exhibit important limitations. They frequently fail to capture the consequences of variants located far from transcription start sites (TSSs) (19), and in some cases predict the wrong direction of effect for alleles linked to expression changes (20, 21). Attempts to address these shortcomings remain limited, as models can be systematically wrong across replicates (22). Moreover, prediction errors are not confined to any single feature dimension, variants distant from a TSS may still be correctly predicted if the model captures sufficiently strong regulatory signals (8, 23). Indeed filtering predictions by predicted effect size, such that only variants predicted to cause large expression changes are retained, remains the most effective strategy for enriching reliable predictions (8, 23). However, this procedure discards a larger proportion of predictions, particularly those in the low-effect regime where most GWAS variants are expected to act (24).

Trustworthiness of individual predictions is a barrier to broader adoption of S2F models in contexts such as translational research or clinical genetics. Moreover, the ongoing debate over whether current training datasets are sufficient to fully resolve cis-regulatory grammar (25, 26) can further erode confidence in S2F predictions. If the underlying grammar remains only partially learned, failures are expected, and knowing which predictions are affected becomes a prerequisite for using these models at all. In structural biology, the transformative utility of AlphaFold (27) was driven not only by its accuracy, but also by its per-residue confidence metrics, which explicitly guide practitioners on which structural regions to interpret and which to set aside. In contrast, regulatory genomics currently lacks an analogous framework for evaluating prediction reliability. While AlphaGenome (10) ranks variant scores against background distributions of common SNPs across scorers and output tracks, an empirical percentile merely reflects where a prediction falls relative to other predicted effects, not whether the prediction is correct. Realizing the full potential of S2F models for disease variant prioritization ultimately requires moving beyond empirical score rescaling toward per-prediction reliability estimation.

We reasoned that whether a variant effect prediction from any S2F model is correct is itself predictable. The variant, the target gene, the tissue context, and the model’s own behavior at a given locus each carry distinct signals indicating whether the model has resolved the local cis-regulatory grammar. To this end, we developed gRely, a meta-modeling framework that assigns each variant–gene–tissue prediction a score of correctness, trained on 1,161 multimodal features against fine-mapped GTEx eQTLs. Applied to Borzoi, gRely achieves 97% sign concordance within its top 20% of predictions across held-out variants, genes, and tissues. Crucially, gRely remains discriminative among low-magnitude variants that standard effect-size filtering discards entirely. Feature attribution provides interpretable insight into systematic base-model failure modes, revealing the genomic and model factors that govern reliability. Furthermore, the learned reliability score generalizes zero-shot to other models, including Enformer (5) and AlphaGenome (10). By providing per-prediction confidence estimates, gRely moves the field from heuristic effect-size filtering toward principled reliability assessment, facilitating the dependable deployment of S2F models in clinical genetics.

## Results

### gRely estimates the reliability of individual variant effect predictions

Trait-associated regulatory variants act by shifting gene expression, and consequently the phenotype distribution (Fig 1A). However, they do so predominantly through small effects (24), placing them in the low-magnitude tail of the predicted |VEP| distribution (Fig 1B). A cutoff that retains predictions by magnitude therefore removes most GWAS-relevant variants along with the unreliable ones. It also removes correct ones. Magnitude separates correct from incorrect Borzoi predictions only partially: 37% of correct predictions fall below 0.1 (Supplementary Fig S1). A reliability estimate that is informative independent of predicted effect size would avoid this trade-off.

**Figure 1:**
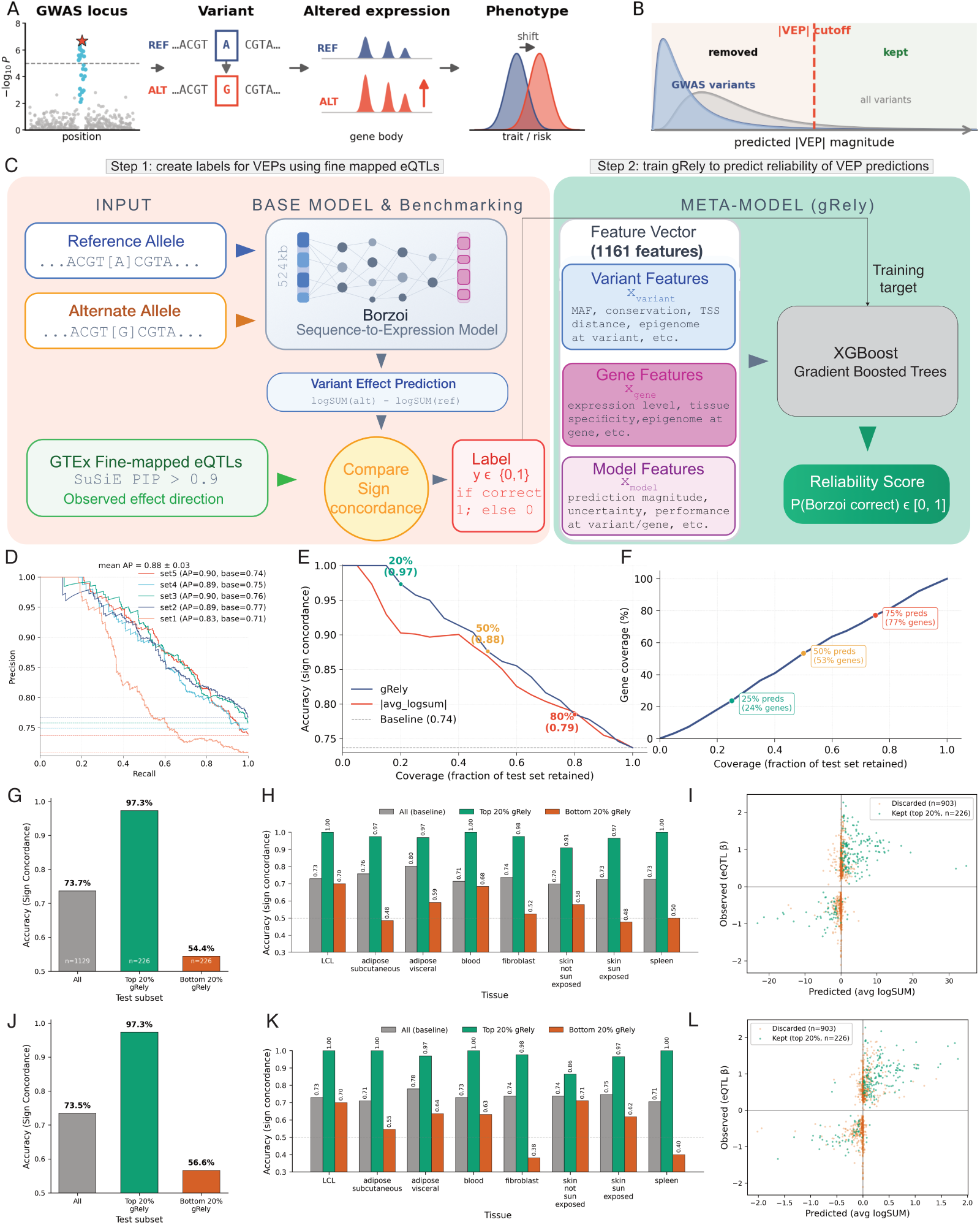
gRely assigns a reliability score to individual VEP and recovers accurate predictions that effect-size filtering discards. **A.** Schematic of how genetic variants alter gene expression to affect trait risk. A GWAS locus shows the lead variant (red star) and associated variants (cyan). The reference allele (blue) and alternate allele (red) produce different expression levels across the gene body, shifting overall disease risk. **B.** Schematic of how filtering by effect size removes most GWAS variants. Distribution of predicted effect magnitudes (|VEP|, x-axis) for all variants (grey) vs. GWAS variants (blue). **C.** Overview of gRely. (Step 1) We label each (variant, gene, tissue) tuple by Borzoi VEP sign concordance against fine-mapped GTEx eQTLs; (Step 2) we train an XGBoost meta-model on 1,161 variant/gene/model features to predict *P* (Borzoi correct). **D.** Precision (y-axis) vs. recall (x-axis) on the sign concordance task for five held-out tissue-set folds (colours); dashed lines, per-fold random baselines. **(E)** Sign concordance accuracy (y-axis) across coverage, the fraction of the test set retained (x-axis), for gRely (blue) and |avg logSUM| thresholding (red); dashed line, unfiltered baseline. **(F)** Percentage of unique genes retained (y-axis) at each prediction-level coverage threshold (x-axis). **(G–I)** Borzoi performance after gRely filtering. **(G)** Sign concordance (y-axis) for all predictions on set5 (*n* = 1,129), the top 20% and the bottom 20% by gRely score (*n* = 226 each; x-axis). **(H)** The same three subsets (colours) within each held-out tissue (x-axis). **(I)** Observed eQTL effect size (*β*; y-axis) vs. predicted VEP (avg logSUM; x-axis) for predictions retained in the top 20% by gRely (green, *n* = 226) and discarded (orange, *n* = 903). **(J–L)** As in **(G–I)**, with gRely applied to AlphaGenome predictions.

We apply gRely to Borzoi VEPs computed for GTEx v8 (28) eQTLs. Each data point is a Borzoi VEP for a (variant, gene, target tissue) tuple. We label each tuple by success in sign concordance task: the label is 1 if the direction of the Borzoi VEP matches the direction of the fine-mapped eQTL effect, and 0 otherwise (SuSiE PIP > 0.9; Fig 1C, Step 1). We featurized each tuple using 1,161 multimodal descriptors (**Methods**) spanning four complementary domains: (i) variant-level context (e.g., local chromatin accessibility centered on the variant locus), (ii) gene-level properties (e.g., baseline expression in training data), (iii) tissue context (e.g., tissue specificity), and (iv) base-model behavior (e.g., cross-replicate agreement). Crucially, to ensure out-of-distribution generalization, we do not encode gene or tissue identity as categorical variables. Instead, genes and tissues are parameterized entirely by continuous functional properties, enabling gRely to score any gene and tissue. Using this feature representation, we trained an XGBoost meta-model (29) that outputs a continuous reliability score between [0, 1] for any candidate prediction (Fig 1C, Step 2).

We evaluated gRely by holding out tissues and chromosomes jointly. We partitioned GTEx tissues into five disjoint sets, and within each fold trained on chr4–chrX and tested on chr1–chr3 (**Methods**). No gene, variant, or tissue in a test set appears in the corresponding training set. gRely separates correct from incorrect Borzoi VEPs on held-out data, with a mean average precision of 0.88 ± 0.03 across folds (Fig 1D).

### gRely outperforms effect-size filtering at matched coverage

Filtering by predicted effect magnitude is the standard approach to enriching reliable VEPs (8, 23). We compared the two criteria directly by ranking held-out predictions either by gRely score or by |avg logSUM|, and measuring sign concordance against the fraction of retained predictions (Fig 1E, Supplementary Fig S2). gRely exceeds magnitude filtering at every coverage below 50%. At 20% coverage gRely reaches 0.97 sign concordance against 0.91 for magnitude filtering, from an unfiltered baseline of 0.75. Above 50% coverage the two criteria converge, as retaining most predictions leaves little room for either to discriminate.

We next examined the predictions gRely retains and discards at a fixed threshold. Filtering to the top 20% of predictions by reliability score raises sign concordance from 73.7% to 97.3% (Fig 1G). The bottom 20% falls to 54.4%, close to chance. gRely therefore identifies unreliable predictions as well as reliable ones. The separation holds in every tissue tested (Fig 1H). Baseline sign concordance ranges from 0.70 to 0.80 across the eight tissues; the top 20% by gRely score ranges from 0.91 to 1.00, and the bottom 20% from 0.48 to 0.70. We see similar pattern across multiple held out tissue folds (Supplementary Table S1, Supplementary Fig S3).

Retained predictions span the full range of predicted effect magnitudes, including variants with |avg logSUM| near zero that a magnitude cutoff would discard (Fig 1I). Binning the test set by predicted magnitude shows the size of this effect (Supplementary Table S2, Table 1). At 50% coverage, magnitude ranking retains nothing below |avg logSUM| of 0.1. gRely retains 273 of these 1,653 predictions and raises their sign concordance from 0.616 to 0.729. In the 0.1–0.2 bin it retains 109 of 356 and raises concordance from 0.677 to 0.807. gRely improves accuracy in the low magnitude bins, so the score carries information that magnitude does not. The same pattern is present in each fold, with the low-magnitude bins showing the largest gain (Supplementary Fig S4).

**Table 1:**
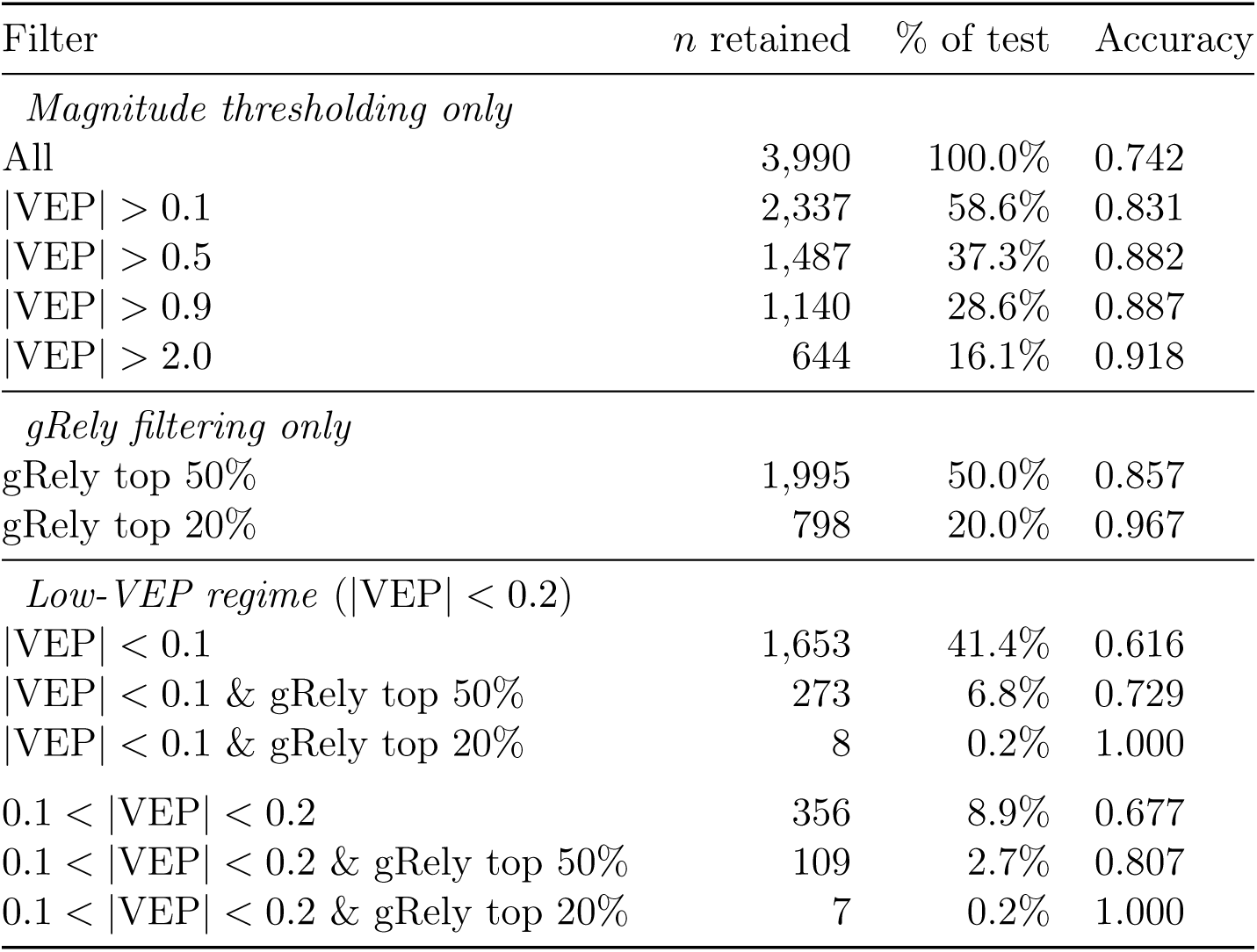
Sign concordance under VEP magnitude thresholding and gRely quantile filtering. Predictions are pooled across the five tissue-set folds (*n* = 3,990 in total). Magnitude thresholding uses a fixed |avg_logSum| cutoff and gRely rows retain the top-scoring quantile. The final block restricts to the low-VEP band that magnitude thresholding discards, and shows that gRely recovers accurate predictions within it.

S2F models can also assign a large effect magnitude to a variant while predicting the wrong direction (21), and these predictions survive magnitude filtering by construction. gRely was able to discard those predictions as well (Fig 1I, Supplementary Fig S3).

### gRely identifies reliable genes and reliable variants within them

Some genes are predicted well by Borzoi and some are not. We asked whether gRely picks this up. We took the genes containing the top 20% of predictions and scored every variant assigned to those genes, including variants gRely did not retain (Supplementary Fig S5). These genes are 16–18% of genes in a fold, and sign concordance over all of their variants is 0.92–0.98, against per-fold baselines of 0.71–0.77. The genes containing the bottom 20% of predictions are 22–26% of genes, and sign concordance over all of their variants is 0.58–0.62. Borzoi is reliable for some genes and unreliable for others, and gRely separates the two.

Selecting accurate genes does not immediately mean selecting few genes. Gene coverage tracks prediction coverage almost exactly: retaining 25%, 50%, and 75% of predictions retains 24%, 53%, and 77% of unique genes respectively (Fig 1F, similar patterns in other folds Supplementary Fig S6). A filter that worked by collapsing onto a handful of well-predicted loci would retain far fewer genes at the same coverage.

If gRely only picked out good genes, every variant in a retained gene would be as reliable as the ones gRely kept. We tested this by repeating the expansion at every threshold rather than at 20% (Supplementary Fig S7). Adding the discarded variants of a retained gene back in lowers sign concordance at every coverage below 100%. The variants gRely keeps are more reliable than the gene they belong to. gRely ranks variants within a gene, not only genes against each other.

### gRely estimates transfer to other S2F models

A meta-model can be built for any base model. However, it requires VEPs and model-derived features for every tuple in the training set, which can be resource intensive in terms of storage and compute. We therefore asked whether the scores we already have carry over. We recomputed nothing. For each (variant, gene, tissue) tuple we kept the reliability score gRely produced from Borzoi features, and asked whether it predicts the correctness of a different model’s VEP on that same tuple. We applied the Borzoi-trained gRely model to AlphaGenome (10) and Enformer (5) VEPs.

For AlphaGenome (10), the top 20% of predictions by reliability score reach 97.3% sign concordance, from an unfiltered baseline of 73.5%, and the bottom 20% falls to 56.6% (Fig 1J). The per-tissue performance was 0.86–1.00 in the top 20% and 0.38–0.71 in the bottom 20% (Fig 1K). Retained predictions again span the low-magnitude regime (Fig 1L). The other four folds behave the same way (Supplementary Fig S8).

Enformer (5) was the harder test. Its unfiltered sign concordance is 59.5% to 67.3% across the five folds, well below Borzoi’s. If gRely worked only because the base model was already accurate, there would be little for it to separate here. It separates Enformer’s predictions in every fold, raising the top 20% to 83.9%–90.3% and lowering the bottom 20% to 45.8%–58.5% (Supplementary Fig S9). gRely does not need an accurate base model to be useful. The base model still sets the ceiling: Enformer’s top 20% ends 8–11 points below Borzoi’s.

Scores derived entirely from Borzoi predict where two other models are correct. The three architectures are not failing independently. They share the variant–gene pairs they get right and the ones they get wrong, which is why a single set of reliability scores works for all three. This means agreement between S2F models is not evidence that a prediction is correct (as seen in (22)), so ensembling across models will not remove these errors. And a new architecture trained on the same data should be expected to fail on the same variants, which points to the training data or the task rather than to model capacity as the limit (25, 26).

### Feature attribution analysis identifies drivers of prediction reliability

We next asked what features gRely use to identify reliable predictions that transfer across several S2F models. We computed SHAP values (30) for every held-out prediction, applying each fold’s model to its own test predictions only, and pooled the attributions (*n* = 3,990 predictions × 1,161 features; Fig 2A). No attribution comes from data the model saw during training.

**Figure 2:**
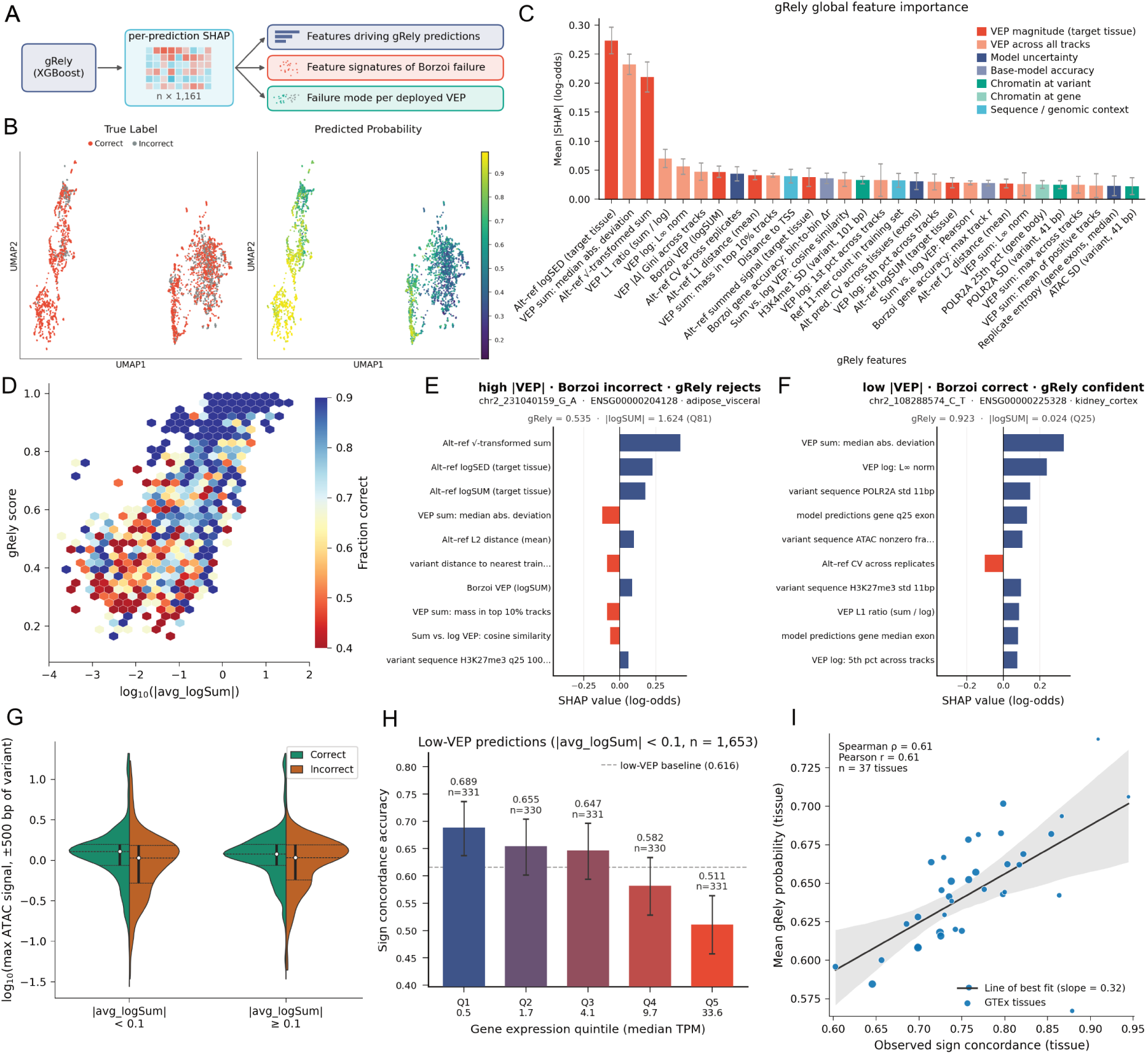
SHAP attribution shows the features gRely uses to separate correct from incorrect predictions. All panels computed on the pooled held-out test set (*n* = 3,990). **A.** Schematic of the interpretation workflow. Per-prediction SHAP values (*n ×* 1,161) are computed from the trained gRely model on held-out predictions and used to rank global feature importance, identify feature signatures of Borzoi failure, and assign a failure mode to individual deployed VEPs. **B.** UMAP embedding of feature attributions, colored by Borzoi sign concordance (left; correct, red; incorrect, grey) and by gRely score (right; color scale). **C.** Global feature importance (mean |SHAP|, log-odds; y-axis) for the highest-ranked features (x-axis), colored by feature type (colors). Error bars show the spread of importance across the five tissue-set folds. **D.** gRely score tracks accuracy across the full range of VEP magnitude. Hexagonal bins of held-out predictions, with gRely score (y-axis) against log_10_(|avg logSUM|) (x-axis), colored by the fraction of predictions in each bin with sign-concordant Borzoi VEP (color scale). Bins containing fewer than 3 predictions are not shown. **E, F.** Per-prediction SHAP attributions for two individual variant–gene–tissue predictions, showing the ten features with largest |SHAP| (y-axes). SHAP value in log-odds (x-axes). (E) A high-magnitude VEP that Borzoi predicts incorrectly and gRely assigns low confidence (gRely = 0.535; |logSUM| = 1.624, 81st percentile) and (F) a low-magnitude VEP that Borzoi predicts correctly and gRely assigns high confidence (gRely = 0.923; |logSUM| = 0.024, 25th percentile). **G.** Distribution of log_10_ maximum ATAC signal within ±500 bp of the variant (y-axis) for correct (green) and incorrect (orange) Borzoi predictions, stratified by VEP magnitude (|avg logSUM| < 0.1 and ≥ 0.1; x-axis). **H.** Sign concordance accuracy declines with target gene expression in the low-magnitude regime. Accuracy (y-axis) for low-VEP predictions (|avg logSUM| < 0.1; *n* = 1,653) binned into quintiles of median TPM of the target gene in the target tissue (x-axis). **I.** Mean gRely score per GTEx tissue (y-axis) against observed sign concordance in that tissue (x-axis); each point is one tissue (*n* = 37).

We can see clear separation in the attribution space. A UMAP embedding splits into distinct regions differing in sign concordance, and gRely scores follow the same split (Fig 2B). Clustering the embedding (Supplementary Fig S10) gives 14 clusters with mean scores from 0.96 to 0.39. As expected, model prediction related features stay dominant, positive in high-scoring clusters and negative in low-scoring ones. However, there are clusters that are driven instead by features such as TSS distance, POLR2A signal at the gene, H3K9ac variability at the variant, etc. Clusters differ in which features they draw on. Top features attributed overlap partially and no two clusters share the same set (Supplementary Fig S10).

The global ranking of feature attributions mirrors these cluster-level findings (Fig 2C). Three features carry most of the weight, and all three describe the magnitude of the predicted effect. Everything below them drops by a factor of three or more and forms a flat tail, where the other feature types appear: replicate disagreement, Borzoi’s accuracy on the target gene, chromatin at the variant and at the gene, and genomic context. Thus, while predicted effect size serves as the strongest individual signal, the remainder is distributed across numerous individually subtle contributions from engineered features.

### gRely uses signals distinct from magnitude

The distribution of feature importance raises the question of whether gRely simply acts as an indirect filter for effect size. We see the two are correlated (Spearman *ρ* = 0.76; Fig 2D), but within a single magnitude region the score varies a wide range. Analyzing per-prediction attributions show what produces it. For a high-magnitude variant that Borzoi predicts with the wrong sign, replicate dispersion, distance to the nearest training region, and disagreement between the sum and log scoring functions pull the score down to 0.535 (Fig 2E). For a low-magnitude variant that Borzoi predicts correctly, POLR2A and ATAC signal at the variant raise it to 0.923 (Fig 2F). Neither call follows from the magnitude.

No secondary feature is usable on its own. Gene expression and replicate disagreement features are ranked highly by gRely yet track the score loosely (Supplementary Fig S11; *ρ* = −0.17 and +0.13), with concordance varying along the score axis rather than the feature axis. Replicate consistency, chromatin at the variant, and concentration of the predicted signal across tracks all differ between correct and incorrect predictions within both the |avg logSUM| < 0.1 and ≥ 0.1 strata (Fig 2G; Supplementary Fig S12), so the separation is not magnitude in another form. The distributions overlap substantially in every case. gRely aggregates them.

### Reliability varies within the regimes where magnitude and distance filters offer no guidance

Effect-size magnitude and TSS distance are the two filters currently used to enrich reliable VEPs. We asked whether reliability still varies within the strata each filter cannot separate.

S2F models can rank the magnitude of a variant’s effect correctly while predicting the wrong sign (21, 22). We asked what distinguishes these direction errors. We labeled each held-out prediction by sign concordance and by magnitude concordance, and restricted to predictions Borzoi ranks correctly in magnitude. We compared these direction errors to sign-correct predictions matched on |VEP| at the variant level, so that the two groups differ in direction alone (*n* = 78 and 74; Supplementary Fig S13). We then screened all the features for separation between them. Twenty-eight of the 30 most discriminative features belong to the target gene expression family, which comprises 109 of the 1,161 features. Direction errors occur at more highly expressed target genes; Supplementary Fig S13). gRely’s attributions on expression-family features track target gene expression and higher expression lowers the predicted reliability. At matched |VEP|, the expression channel separates direction errors from sign-correct predictions, while neither the magnitude channel nor the full attribution does (Supplementary Fig S13).

The expression axis is not confined to direction errors. It also degrades accuracy in the low-magnitude regime, where magnitude filtering offers no guidance at all: correct low-VEP predictions fall on genes with lower observed expression than incorrect ones (median 3.5 vs 5.4 TPM), and accuracy declines monotonically across expression quintiles, from 68.9% for the lowest-expressed genes to 51.1% for the highest (Fig 2H). The same vulnerability appears in enhancer–gene benchmarking, where ENCODE-rE2G performance was lower for housekeeping genes than for other genes (31). Highly expressed genes are therefore a systematic blind spot of current models.

Distance to the target TSS is the other axis along which S2F failures are known to concentrate (19). Ranking held-out predictions by gRely score retains more sign-concordant predictions than ranking by TSS proximity or by |VEP| at any stringent threshold (0.97 vs 0.92 and 0.91 at 20% retention; Supplementary Fig S14). Sign concordance accuracy falls with distance as expected. Distance, however, is a poor proxy for reliability at the level of an individual prediction. Within every distance stratum tested, the top quintile of gRely scores is far more concordant than the bottom quintile, and predictions within 1 kb of the TSS that fall in the bottom quintile are *less* concordant (0.64) than the unfiltered set as a whole (0.74): proximity to the TSS does not make a prediction safe. Conversely, predictions 10–100 kb from the TSS that gRely scores in the top quintile reach 0.97 concordance, recovering reliable predictions in exactly the regime a distance cutoff would discard. gRely does encode distance in aggregate as 37% of variants within 1 kb fall in the global top quintile versus 12% at 10–100 kb. However, it does not reduce to it (Supplementary Fig S14).

### gRely estimates reliability for tissues without eQTL benchmarks

The promise of S2F models is that they can nominate causal variants and prioritise candidates for experimental follow-up. Currently, to know whether a prediction can be trusted on a specific tissue, we use fine-mapped eQTLs for the tissue in question to report model performance. However, most cell types of interest have none, while Borzoi predicts tracks for them. gRely fills this gap by estimating model reliability for unseen tissues.

We do not hard code tissue identity, rather through quantities computed from its matching Borzoi track: observed expression of the target gene in that tissue, Borzoi’s accuracy on that track, the predicted signal over the gene in that track, etc. All are available for any tissue Borzoi has a track for. Our fold assignment holds out whole tissue sets, so every prediction we score comes from a tissue the meta-model never saw. Across the 37 GTEx tissues, mean gRely score ranks tissues by their observed sign concordance (Spearman *ρ* = 0.61; Fig 2I).

Beyond just filtering for reliable predictions, gRely supports inspection of individual predictions (Supplementary Fig S15), reporting the features such as chromatin context at the variant, that context relative to all other variants, and the per-prediction SHAP attribution summed by feature category. In the example shown, Borzoi predicts the wrong direction of effect for a variant only 551 bp from the target TSS. gRely assigns it a sign-concordance reliability of 0.31, and the attribution shows the low score is driven by the local chromatin context, specifically H3K9me3 at the variant, rather than by the magnitude of the predicted effect or Borzoi’s accuracy on the gene. Consistent with this, the variant lies in repressed chromatin: H3K9me3 and H3K27me3 exceed the mean across held-out variants while accessibility and active marks fall below it. gRely thus serves not only to prioritize reliable predictions but also to identify the contextual features underlying individual S2F failures.

## Discussion

S2F models suffer from several known failure modes, including poor performance on distal variants (19) and frequent disagreement among independently trained replicates (22). To address this, we developed gRely, a meta-model that comprehensively aggregates features describing the variant, target gene, tissue context, and the base model’s own behavior to estimate whether an individual VEP can be trusted. gRely recovers accurate predictions from the low-|VEP| region that magnitude thresholding discards. Importantly, it ranks variants within a single gene rather than merely distinguishing well-predicted genes from poorly predicted ones. It introduces individual prediction trustworthiness (32) to regulatory genomics, a much-needed filter for prioritizing causal variants.

Remarkably, a meta-model trained entirely on Borzoi (7) predictions transfers directly to Al-phaGenome (10) and Enformer (5) without retraining. Despite differences in model size, receptive field, and output tracks, these architectures succeed and fail on similar variant–gene–tissue tuples. This shared vulnerability implies that prediction reliability is fundamentally a property of the biological context and the underlying training data, rather than model capacity. Consequently, mere agreement between distinct architectures is not evidence of correctness; simple ensembling cannot resolve these errors. If new architectures are trained on the same data, they will likely fail in the same places, suggesting that resolving these blind spots will require rethinking the training tasks and data composition themselves (25, 33).

The scale of these unreliable predictions is alarming. The bottom quintile of Borzoi unreliable predictions achieves 54% sign concordance, which is essentially random chance; and the bottom half reaches only 63%. Without a reliability estimate, these predictions will remain indistinguishable from genuine biological signal, severely confounding both GWAS prioritization and clinical variant interpretation. The problem grows as these models are deployed at scale. Predictions can be precomputed for every possible substitution in the genome, so the task is no longer running a model but deciding which of billions of scores deserve an experiment. The heuristics available for that decision are blunt. Filtering by effect size discards the low-effect variants that most complex traits act through, and filtering by proximity to the TSS fares no better: near-TSS predictions in the bottom gRely quintile are less concordant than the unfiltered set as a whole, while distal predictions in the top quintile reach 97%. Neither heuristic separates predictions well, which leaves reliability estimation as the practical solution. We propose that future S2F architectures should intrinsically incorporate a reliability head trained jointly with the core predictor. Until they become standard, external meta-models like gRely will be critical for model deployment.

Feature attribution also highlights specific biological regimes where current models systematically break down. For instance, direction errors, where a model predicts the correct VEP magnitude but the wrong sign, are concentrated at highly expressed target genes. Beyond baseline expression, our clustering of failure modes reveals distinct error regimes that are dominated by chromatin marks at the variant locus. These features provide a clear roadmap for the next generation of models. Given gRely scores transfer across models, merely scaling model capacity is unlikely to resolve these shared blind spots. Instead, changing the information available to the model is one of the few levers expected to meaningfully improve accuracy. Architectures designed to condition on observed functional data alongside DNA sequence at inference, such as Nona (34), can utilize these signals directly. Consequently, the highly ranked functional tracks identified by gRely provide a principled starting list of which contextual features these new models should condition on to avoid the pitfalls of sequence-only prediction.

Crucially, gRely remains effective where fine-mapped eQTL ground truth is absent. Because tissues are parameterized through continuous features derived from their matching functional tracks rather than rigid categorical identifiers, gRely can provide reliability of predictions in tissues it has never seen during training. The mean gRely score accurately ranks held out GTEx tissues by their observed sign concordance (*ρ* = 0.61). At the individual prediction level, these scores are coupled with feature attributions, transforming a reliability metric from a simple black-box verdict into an interpretable diagnostic that isolates the specific drivers, such as unfavorable local chromatin context or underlying replicate disagreement, causing the model’s failure in a specific tissue.

Several limitations remain in our framework. While fine-mapped eQTLs represent the most confident catalog of causal regulatory variants currently available at genome scale (35–37), they remain a proxy for ground truth. Consequently, gRely estimates whether a prediction agrees with a fine-mapped eQTL rather than providing an absolute measure of biological correctness, inherently adopting the biases and coverage limitations of the underlying catalog. For instance, our minor allele frequency (MAF) cutoff of 0.01 excludes rare variants, and several GTEx tissues were omitted entirely due to insufficient fine-mapping power. Additionally, our chromatin features are averaged across cell types, which restricts the tissue-specific context available to the meta-model. Moreover, our hand-engineered features likely miss complex, high-dimensional signals that direct access to the base model’s latent representations or base-pair-level attributions might reveal. Future work should explore richer, more comprehensive featurization strategies to capture these nuances.

## Methods

### Base model

We used Borzoi (7), a S2F model that takes a 524,288 bp DNA sequence as input and outputs predicted coverage for RNA-seq, CAGE, DNase-seq, ATAC-seq, and ChIP-seq experiments across 7,611 tracks at 32 bp resolution. During training, loss calculation was restricted to the central 196,608 bp of each input (6,144 bins). We used the four independently trained Borzoi replicates released with the model.

### Variant effect scores

For variant effect prediction, we centered each sequence on the target variant and extracted features from the central 196,608 bp window (6,144 bins); variants with no exon overlapping this window were excluded. For each (variant, gene, target tissue) tuple, we centered the input sequence on the variant and predicted the reference and alternate alleles with each of the four replicates. Within the prediction window, we selected the bins overlapping any exon of the target gene and the GTEx track(s) matching the target tissue (Supplementary Table S3). Predicted coverage was summed over exon bins for each allele and the variant effect was scored as the difference of the log_2_-transformed summed coverage values:

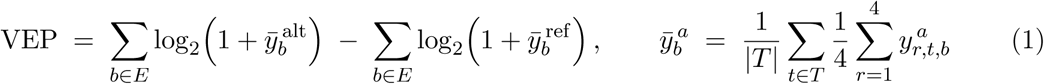

where *y^a^_r,t,b_* is the coverage predicted by replicate *r* in track *t* and bin *b* for allele *a ∈ {*ref, alt*}*, *T* is the set of Borzoi GTEx track(s) matching the target tissue (Supplementary Table S3), and *E* is the set of bins overlapping any exon of the target gene, restricted to the central 6,144 bins of the output window. Both averages are taken before the log transform, so a single score is computed per tuple rather than one per replicate.

Scores were averaged over the four replicates. Throughout the paper, VEP and logSum denote this replicate-averaged score in the target tissue, and |VEP| its absolute value. Tuples whose target gene had no exon within the prediction window were excluded; every scored gene therefore has an exon within 98,304 bp of the variant.

### Ground truth and labels

We used SuSiE (35, 36) fine-mapped eQTL credible sets from the eQTL Catalogue (37) as proxy ground truth to train and evaluate our reliability framework. Among its outputs, Borzoi makes predictions for 89 GTEx tracks, each corresponding to an individual sample from a human tissue. These tracks span 41 tissues or conditions in GTEx. Four tissues with insufficient sample sizes for SuSiE fine-mapping were excluded, leaving a final dataset covering 37 GTEx cell types.

To curate our variant set, we applied the following filters:

- PIP threshold: retain variants with SuSiE PIP > 0.9.
- Variant type: restrict to single-nucleotide variants (SNVs).
- Blacklist exclusion: remove variants overlapping hg38 blacklist regions (v2) (38).
- Frequency filter: require minor allele frequency (MAF) > 0.01 using GTEx summary statistics.

Each candidate data point was defined as a (variant, gene, target tissue) tuple, where the variant had been fine-mapped to the gene in a specific cell type. For each tuple, Borzoi predictions were obtained by centering input sequences on the variant, extracting exonic bins within the model’s central 6,144-bin output window, and computing a logSUM score (as described in borzoi (7)) between reference and alternative alleles. Predictions were generated using four independently trained Borzoi replicates, and the average prediction across replicates was used. Only the output track(s) corresponding to the target tissue was considered, as specified in Supplementary Table S3.

Each data point was labeled by comparing Borzoi’s predicted direction of effect against the fine-mapped eQTL: label = 1 if the predicted sign matches the eQTL direction (sign-concordant), label = 0 otherwise. This sign concordance label served as the binary target for gRely.

For each labeled instance, we extracted variant-level, gene-level, and model-derived features as inputs to gRely. Importantly, all features provided to gRely can be computed for any variant–gene pair in the genome, as long as the target gene contains exons within 98,304 bp (half of the 196,608 bp central prediction window used during Borzoi training). To prevent information leakage, we excluded features derived directly from eQTL summary statistics (fine-mapping PIP, credible set metrics, eQTL p-value and standard error, allele frequency, and allele counts). The evaluation followed a 5-fold cross-validation scheme in which each fold holds out one tissue group (sets 1–5; Table 2) and chromosomes 1–3, with training on the remaining tissues and chromosomes 4–X; this joint chromosomal–tissue split ensures that neither the variants nor the tissue contexts seen at test time appear during training.

**Table 2:** Composition of the five held-out tissue sets. The 37 GTEx tissues were partitioned into five disjoint sets grouped by organ system. In each cross-validation fold, one set is held out together with chromosomes 1–3, and the meta-model is trained on the remaining four sets on chromosomes 4–X. Tissue names are given in GTEx display form; the corresponding eQTL Catalogue identifiers are listed in Supplementary Table S3. The last column gives the number of held-out test predictions in that fold (total *n* = 3,990).

| Set | Grouping | Tissues | Members | Test $n$ |
| --- | --- | --- | --- | --- |
| 1 | Neural and endocrine | 7 | Brain (amygdala), Brain (cerebellar hemisphere), Brain (cortex), Nerve (tibial), Pituitary, Adrenal gland, Thyroid | 838 |
| 2 | Cardiovascular, muscle, lung, kidney | 7 | Artery (aorta), Artery (tibial), Heart (atrial appendage), Heart (left ventricle), Muscle (skeletal), Lung, Kidney (cortex) | 778 |
| 3 | Digestive | 8 | Esophagus (gastroesophageal junction), Esophagus (mucosa), Stomach, Small intestine, Colon (sigmoid), Colon (transverse), Liver, Pancreas | 719 |
| 4 | Reproductive and glandular | 7 | Ovary, Testis, Uterus, Vagina, Prostate, Breast (mammary tissue), Minor salivary gland | 526 |
| 5 | Blood, immune, skin, adipose | 8 | Whole blood, Lymphoblastoid cell line, Spleen, Skin (not sun exposed), Skin (sun exposed), Cultured fibroblasts, Adipose (subcutaneous), Adipose (visceral) | 1,129 |

### gRely meta model

We formulated gRely as a supervised binary classification task, where the goal was to estimate a score for Borzoi’s prediction for a given (variant, gene, tissue) tuple was correct. We implemented gRely using gradient-boosted decision trees (XGBoost). The model was trained with the logloss objective and the following hyperparameters: 300 estimators, maximum tree depth of 5, learning rate of 0.05, subsample ratio of 0.8, column subsample ratio (colsample_bytree) of 0.8, and random seed of 42. To address class imbalance, we set scale_pos_weight to the ratio of negative to positive samples in the training set. The model’s output was interpreted as a score for Borzoi’s direction of effect prediction is correct for a given (variant, gene, tissue) tuple.

### Gene Annotations

Gene annotations were obtained from GENCODE v41 (39). For each gene, we considered all annotated isoforms and defined the gene’s target interval as the union of all exon start and end sites across isoforms. This ensured that exonic regions from every transcript variant were included in downstream analyses. Transcription start sites (TSSs) were derived using the transcript annotation provided in GENCODE.

### gRely features

Each tuple was represented by 1,161 features spanning four domains: the variant, the target gene, the target tissue and the base model’s own behaviour. Two constraints governed feature design. First, every feature is computable for any variant–gene–tissue combination in which the gene has an exon within the Borzoi prediction window, so that gRely can score variants. Second, no feature encodes the identity of a gene, tissue or variant categorically; genes and tissues enter only through continuous functional descriptors, which makes held-out genes and tissues predictable at test time. We excluded all quantities derived from eQTL summary statistics (fine-mapping PIP, credible-set metrics, p-values, standard errors, allele frequencies and allele counts). All 1,161 features were used; no feature selection was performed. Missing values were left as NaN and handled natively by XGBoost.

### Gene related features

**Sequence–chromatin features** For each gene body, we computed GC content and summary statistics from a panel of chromatin-associated signal tracks derived from EpiMap ‘Other’ group (40), including ATAC-seq, DNase-seq, histone modifications (H3K27ac, H3K4me1, H3K4me3, H3K9ac, H3K9me3, H3K27me3, H3K36me3), CTCF binding, and RNA polymerase II occupancy (POLR2A). The tracks were created by averaging across multiple cell types. For each track, we calculated the mean, median, maximum, minimum, standard deviation, 25th and 75^th^ percentiles, fraction of non-zero positions, fraction of missing values, and total signal across the gene span.

**Model performance on the gene** We evaluated Borzoi’s predictive accuracy at the gene level using the Pearson correlation between predicted and observed targets within the centered 6,144-bin slice of the model’s full 16,384-bin output. To select the target data for each gene, we considered all TFRecord intervals overlapping the gene’s genomic interval and selected the interval with the maximum overlap length; ties were broken by choosing the interval whose center was closest to the gene center. This yields the data point with maximal overlap (and closest center if needed) for feature extraction. For each gene and each of the four trained model replicates, we computed performance metrics at both the track and bin levels, and then averaged results across replicates.

**Per-track correlations across sequence bins** For each track, correlations were computed across the 7,611 sequence bins overlapping the gene. After averaging across replicates, we summarized performance by: mean, standard deviation, minimum, maximum, median, 90th percentile, and the fraction of correlations exceeding 0.8.

**Per-bin correlations across tracks** For each genomic bin, correlations were computed across the 6,144 tracks. After averaging across replicates, we summarized performance by: mean, standard deviation, minimum, maximum, median, 90th percentile, fraction of correlations above 0.8, and additionally the mean and standard deviation of first differences along the sequence.

**Cell type–specific performance** For the GTEx target tissue corresponding to each (variant, gene) pair, we identified the matching Borzoi track. We reported the correlation for this track, its deviation from the GTEx-wide mean and from the mean of all other tissues, and a z-score relative to the full distribution of GTEx track correlations.

**Observed Gene Expression Features** We derived gene-level features from Borzoi’s per–cell-type × genomic-bin target matrices (89 GTEx tracks) available in TFRecords with 6,144 bins spanning a 196,608 bp output window. For each gene, the TFRecord was selected as described above. Features were computed both globally (over all bins) and within exon-overlapping bins.

**Global features** Distributional statistics: sum, mean, median, minimum, maximum, standard deviation, skewness, kurtosis, 25th and 75th percentiles, Gini coefficient, and tissue-specificity index (*τ*).

Across-cell variability (per bin): variance and range across cell types for each bin, summarized by the mean and maximum across bins.

Across-bin variability (per cell type): variance across bins for each cell type, summarized by the mean and maximum across cell types.

Entropy: Shannon entropy per bin (after normalizing across cell types), summarized by mean, median, and maximum.

Sparsity and peakiness: fraction of entries < 1e-3 and the number of entries above the 95th percentile of the matrix distribution.

**Cell type–specific features** For each cell type, we computed mean, sum, maximum, standard deviation, and sparsity across bins, both globally and within exons. These were further summarized across cell types:

Cross-cell variability: coefficient of variation, *τ*, Gini coefficient, and the top-one fraction (ratio of maximum to total).

Specificity relative to other cell types: z-score and fold-change relative to the mean of all other cell types, fraction of total expression attributable to the cell type, and a ratio-based specificity index defined as 1–mean(relative expression of others).

For each gRely data point - defined by a (variant, gene, target tissue) tuple - we used all global features and from cell type specific features only the features corresponding to the target tissue.

### Variant related features

**Sequence–chromatin features** Sequence-derived features included GC content within multiple centered windows (11, 21, 41, 101, and 1001 bp) and k-mer occurrence counts for 11-, 21-, and 41-mers in the reference and alternate alleles within the training set. Chromatin context was characterized using the same EpiMap (40) chromatin-associated signal tracks used for gene-level features, including ATAC-seq, DNase-seq, histone modifications (H3K27ac, H3K4me1, H3K4me3, H3K9ac, H3K9me3, H3K27me3, H3K36me3), CTCF binding, and RNA polymerase II occupancy (POLR2A). For each track, we computed the mean, median, maximum, minimum, standard deviation, 25th and 75th percentiles, fraction of non-zero positions, fraction of missing values, and total signal within each window.

**Conservation features** Evolutionary conservation was quantified using phyloP scores (41, 42) at the variant position and averaged over 100 bp and 1,000 bp windows centered on the variant.

**Genomic context features** We computed distances from the variant to the transcription start site (TSS) of the target gene, the nearest centromere, and the nearest telomere. Additional features described the variant’s spatial relationship to training regions, including whether it overlapped any training region and the distance to the nearest training region center.

**Local eQTL density features** For each candidate variant, we quantify positive eQTL density in symmetric windows centered at the variant (±1 bp, ±1 kb, ±10 kb) on the same chromosome. For each window, we compute counts both across all tissues: (i) the number of positive eQTL rows (variant–gene–tissue records) and (ii) the number of unique variants, defined by unique (chrom, pos).

**Model performance on the variant** We evaluated Borzoi’s predictive accuracy at the variant level using per-variant Pearson correlation vectors. To select the target data for each variant, we considered all TFRecord intervals overlapping the variant and selected the interval whose center had least distance to the variant. This yields the data point with least distance from center to variant for feature extraction. Correlations were averaged across the four model replicates before summarization. Variant-level performance was assessed along two axes: across tracks and across sequence bins, with additional evaluation of the mapped GTEx target cell type.

**Across tracks (along sequence; 7,611 tracks)** For each variant, correlations were computed across all tracks. After averaging across replicates, we summarized by: mean, standard deviation, minimum, maximum, median, 90th percentile, and the fraction of tracks with correlation exceeding 0.8.

**Across sequence bins (along track; 6,144 bins)** For each variant, correlations were computed across bins within the 6,144-bin central region. After averaging across replicates, we summarized by: mean, standard deviation, minimum, maximum, median, 90th percentile, fraction of bins with correlation above 0.8, and first-difference statistics along the sequence (mean and standard deviation of adjacent bin-to-bin changes).

**GTEx target cell type performance** For the target GTEx tissue mapped to each variant, we averaged its track(s) and, over the gene’s bins, reported totals of the raw signal, log-transformed signal (log2+1), and square-root-transformed signal, along with the mean log2 of per-bin totals; also reported the corresponding differences between alternate and reference for each of these aggregates.

### Model related features

**Model prediction difference features across 89 GTEx tracks** We used four independently trained model replicates to predict per-track outputs for the reference and alternate alleles, saving only the 89 GTEx tracks. Predictions were cropped to the centered 6,144-bin window. For each allele, we stacked replicates to compute replicate means and per-bin replicate dispersions.

**Replicate aggregation and central cropping** Center the 6,144-bin window; average predictions across replicates per bin and track; compute per-bin replicate dispersion.

**Effect-size across alleles (using replicate-averaged predictions over the center 6144 bins)** L1 distance: Mean across tracks of the total absolute difference across bins. L2 distance: Mean across tracks of the Euclidean norm of differences across bins. Maximum absolute change: Mean across tracks of the signed difference at each track’s bin of maximum absolute change.

**Replicate uncertainty** For reference and alternate alleles separately: mean, maximum, and standard deviation of per-bin replicate standard deviations.

Coefficient of variation (CV): mean over bins of replicate standard deviation / (replicate mean + 1e-6).

Replicate-level delta uncertainty: replicate standard deviation and CV of (alt − ref) predictions.

Agreement fraction: fraction of bins with replicate standard deviation < 0.05 for reference and alternate predictions.

Replicate entropy: mean across tracks of per-bin Shannon entropy across replicates, computed separately for reference and alternate predictions.

**Gene’s exon overlapping bins and target-tissue deltas (using replicate-averaged predictions over the central bins)** Within exon-overlapping bins, we computed per-track means for reference and alternate; and also took their difference per track. We summarize absolute differences across tracks using tissue-specificity index (*τ*), inequality (Gini coefficient), top-one fraction (largest share over total), and coefficient of variation.

For the mapped target tissue, we averaged its track(s) and, over the gene’s bins, reported totals of the raw signal, log-transformed signal (log2+1), and square-root-transformed signal, along with the mean log2 of per-bin totals; also reported the corresponding differences between alternate and reference for each of these aggregates.

**Ref/alt prediction feature bundles on the gene** Using replicate-averaged predictions, we also calculated the ‘Observed Gene Expression Features’ within the gene slice for both reference and alternate alleles.

**Model prediction difference features across all 7611 tracks** Due to the massive storage required to store the predictions, we do not save full predictions on all 7611 tracks. We saved only per-track variant effect scores for four Borzoi replicates. For each replicate, both sumscore and logsum score (length 7,611) were calculated. Replicate predictions were averaged to obtain two final track-level vectors. Features were then computed in three groups:

**Per-vector summaries (applied separately to the sum and log vectors)** Distributional statistics: mean, standard deviation, minimum, maximum, median, and 10th/90th percentiles. Norms: L1 norm (sum of absolute values) and L2 norm. Sign composition: fraction of positive, negative, and zero entries. Threshold exceedances: fraction of entries with absolute values above 0.1, 0.5, and 1.0. Concentration metrics: share of absolute mass in the top-1, top-5, and top-10 entries. Inequality and sparsity: Gini-like coefficient and sparsity proxies over the absolute values.

**Cross-vector relationships** Pearson correlation and cosine similarity between **s**^sum^ and **s**^log^; fraction of tracks with the same sign across both vectors (restricted to tracks where either is non-zero); and ratios of L1 and L2 norms (||**s**^sum^ || _1_/ ||**s**^log^ ||_1_ and ||**s**^sum^ ||_2_/ ||**s**^log^ ||_2_).

**Model-Predicted Gene Features** We derived gene-level features from Borzoi’s replicate-averaged predictions and replicate variability restricted to each gene’s exonic bins intersected with the centered 6,144-bin window.

**Replicate uncertainty (for gene predictions)** We computed summary statistics of replicate dispersion per bin and then aggregated across bins/tracks:

Gene prediction standard deviation: mean, maximum, and standard deviation of per-bin replicate standard deviations.

Gene prediction coefficient of variation: Mean coefficient of variation across bins, defined as replicate dispersion divided by the absolute replicate mean (stabilized).

Gene prediction agreement fraction: fraction of bins with replicate standard deviation < 0.05.

Gene prediction entropy: mean across tracks of the per-bin Shannon entropy over replicate distributions (replicate values shifted non-negative and normalized per bin).

**Target-tissue over the gene slice** After averaging the mapped GTEx track(s), we summarized the distribution over bins:

Level summaries: total predicted signal; total of log2(signal + 1); total of sqrt(signal); and the mean of log2(total signal per bin + 1).

Descriptives: mean, standard deviation, minimum, maximum, median, 90th percentile, fraction of bins with value > 0.8, and the bin position of the maximum.

**Cross-track concentration (means over gene-slice bins)** From per-track means (averaged across the gene’s bins), compute: Tissue-specificity index (*τ*), inequality (Gini coefficient), coefficient of variation, and the top-one fraction (largest track mean divided by the total across tracks).

**Matrix-level summaries within the gene slice** Apply the same “Observed gene expression” summary features to the replicate-averaged prediction matrix, restricted to exon-overlapping bins.

### Tissue related features

The target tissue is never encoded as a categorical variable. It enters only through the subset of the gene-and model-related features above that are evaluated on the Borzoi track(s) matching the tuple’s tissue, rather than pooled across all 89 GTEx tracks, together with the cross-track specificity summaries that place the target tissue relative to the others. Every such quantity is defined for any track Borzoi predicts and none of them requires eQTL data for that tissue, so a tuple from a tissue absent from the training can be scored.

### Evaluation

**Precision–recall** Within each fold, we ranked held-out predictions by gRely score and computed the precision–recall curve against the sign concordance label, summarizing it by average precision. The random baseline for a fold is the fraction of sign-concordant predictions in its test set.

**Coverage–accuracy evaluation and magnitude thresholding** To evaluate the trade-off between prediction confidence and dataset coverage, we swept gRely score thresholds across the held-out test set for each cross-validation fold. For a given fold, we computed gRely predicted scores on the test set (held-out chromosomes 1–3 and held-out tissue set) and defined threshold values corresponding to percentiles of the test-set score distribution. At each threshold *t*, we retained all predictions with gRely score ≥ *t*, and computed: (i) *prediction coverage*, defined as the fraction of test-set predictions retained; and (ii) *sign concordance accuracy*, defined as the fraction of retained predictions where Borzoi’s predicted direction agrees with the eQTL ground truth.

To compare gRely against the standard practice of filtering by predicted effect magnitude, we repeated the same sweep with predictions ranked by |avg_logSum| instead of by gRely score, so that the two criteria are evaluated at matched coverage. However, we note this is not really a fair comparison as the two sets select different sets of variants and magnitude filtering would miss many variants of biological importance. We report sign concordance at fixed cutoffs and at fixed gRely percentiles on the pooled test set.

**Gene coverage** To assess whether gRely filtering preferentially eliminates specific genes, we additionally computed *gene coverage* at each threshold as the fraction of unique genes with at least one retained prediction, relative to the total number of unique genes in the test set.

### Transfer to other sequence-to-function models

gRely scores were computed once from Borzoi-derived features and were not recomputed for other models. To test transfer we obtained VEPs for the same (variant, gene, tissue) tuples from AlphaGenome (10) and Enformer (5), labeled each tuple by whether that model’s VEP sign matched the eQTL effect, and repeated the analyses above, selecting the top and bottom 20% of each fold’s test set by the Borzoi-derived gRely score and reporting sign concordance overall and per tissue. No AlphaGenome-or Enformer-derived features were used and gRely was not retrained.

### AlphaGenome evaluation

AlphaGenome (10) predictions were obtained through the public API for a 1,048,576 bp input sequence centred on the variant, taking the RNA_SEQ output modality (768 tracks) at 128 bp resolution. For each target tissue we used the single AlphaGenome RNA-seq track whose gtex_tissue annotation matches the GTEx tissue of the tuple (Supplementary Table S4). Reference and alternate alleles were predicted separately and the signal was restricted to the output bins overlapping any exon of the target gene, a bin being included if it overlaps an exon by at least 1 bp. The variant effect score was computed as

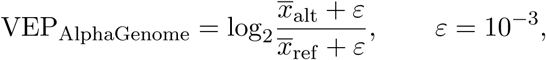

where *x* is the mean predicted signal over the gene’s exonic bins in the target track.

### Enformer evaluation

Enformer (5) is a further test of cross-architecture generalization: it predates Borzoi and predicts 5,313 human tracks at 128 bp resolution over 896 output bins, covering the central 114,688 bp of its input window, four times coarser and roughly half as wide as Borzoi’s 196,608 bp prediction window. We used weights from the TF Hub checkpoint. Reference and alternate alleles were predicted separately for a sequence centred on the variant, placing the output window at ±57,344 bp around it, and predictions were restricted to the output bins overlapping any exon of the target gene under the same any-overlap rule.

Enformer produces no GTEx RNA-seq outputs, so each GTEx tissue was mapped to the closest matching CAGE track (Supplementary Table S5). Five of the 37 tissues are represented by an anatomical, developmental-stage or primary-cell-type proxy rather than a direct match; and three pairs of tissues (sigmoid and transverse colon; gastroesophageal junction and esophageal mucosa; sun-exposed and non-sun-exposed skin) share a single track and therefore receive identical Enformer VEPs for the same variant–gene pair. The variant effect score is calculated using

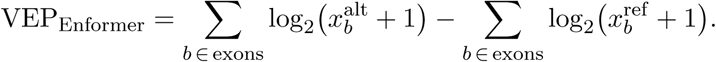

Because Enformer’s output window is narrower than Borzoi’s, a target gene whose exons lie more than 57,344 bp from the variant falls outside it and cannot be scored, although it is within range for Borzoi and AlphaGenome. Such tuples were considered wrong predictions.

### Feature attribution

**SHAP values** We computed SHAP attributions for every held-out prediction using XGBoost’s native implementation, applying each fold’s model to its own test set only. Pooling the five folds gives an *n ×* 1,161 attribution matrix (*n* = 3,990). No attribution derives from data a model saw during training.

**Global feature importance** The importance of a feature is its mean absolute SHAP value over the pooled test set. We report the highest-ranked features coloured by category, with error bars showing the spread of the per-fold mean |SHAP| across the five folds.

**Embedding and clustering** We embedded the pooled attribution matrix with UMAP and clustered it with the Leiden algorithm. For each cluster we report the mean gRely score and the mean attribution of the features ranking in the top 20 by mean |SHAP| within any cluster. Similarity of feature usage between clusters was quantified as the Jaccard index between their top-20 feature sets.

## Data availability

All data used in this study are publicly available. Borzoi model weights were obtained from (https://github.com/calico/borzoi), and the processed data from gs://borzoi-paper/data. Fine-mapped eQTL credible sets used as proxy ground truth were downloaded from the eQTL Catalogue (https://www.ebi.ac.uk/eqtl); the corresponding GTEx v8 summary statistics were obtained from the GTEx portal (https://gtexportal.org). Gene annotations were obtained from GENCODE v41 (https://www.gencodegenes.org). Chromatin signal tracks were obtained from EpiMap (https://compbio.mit.edu/epimap/).

## Code availability

All code used to generate features, train and evaluate gRely is available at https://github.com/cafletezbrant/gRely.

## Acknowledgements

We thank Surag Nair, David Garfield, Xiaoman Xie, Avantika Lal, Laura Gunsalus, and Muhammed Hasan Çelik for helpful discussions and feedback on this work. We also thank Johannes Hingerl, Ayesha Bajwa, Michael Callahan, William Cheney, and Carl de Boer for discussions.

## Author contributions

A.M.R., K.F.B., and G.E. conceived the study, designed the experiments, engineered the features, performed all analyses, and interpreted the results. A.M.R. and K.F.B. wrote the manuscript. K.F.B. supervised the study.

---

**Supplementary Table S1:**
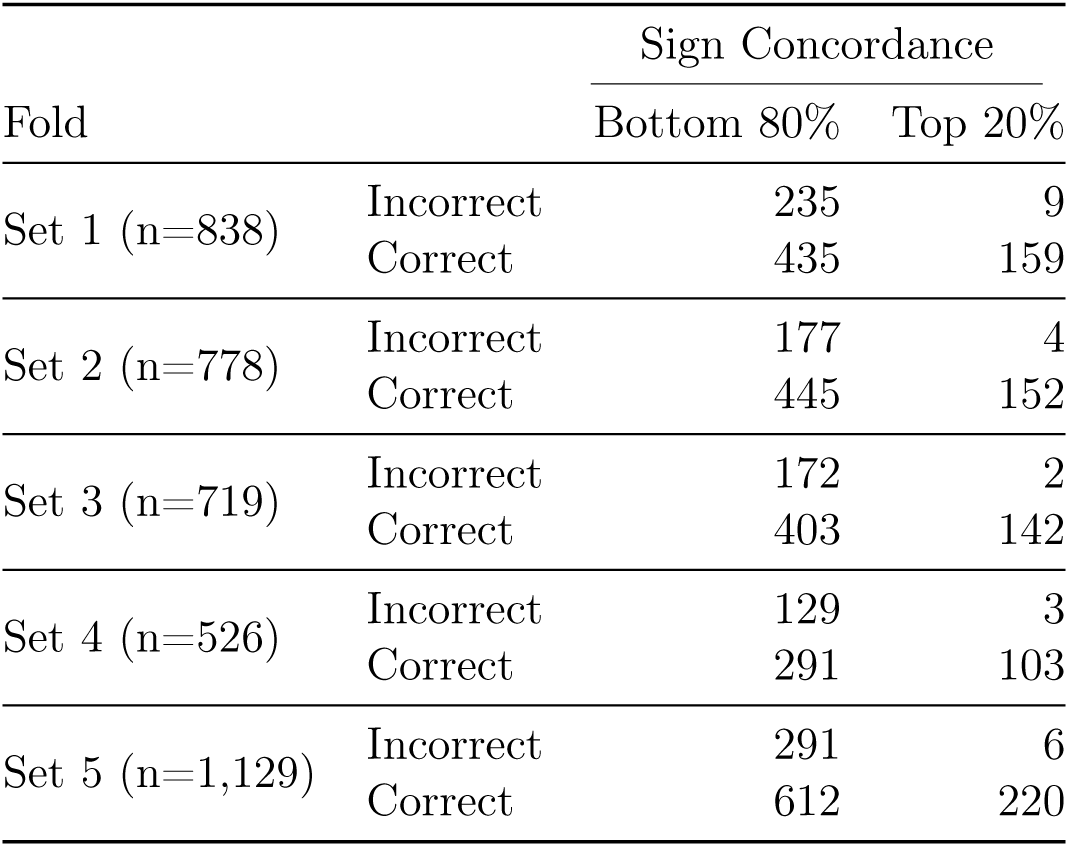
Sign concordance of Borzoi predictions in the top 20% of gRely scores. Within each held-out fold (test chromosomes 1–3, a distinct set of held-out tissues per fold), predictions are ranked by gRely score and split into the bottom 80% and top 20%. The top-20% subset achieves near-perfect sign concordance across all folds (mean 97.0%) while retaining 20% of test-set predictions.

| Fold |  | Sign Concordance |  |
| --- | --- | --- | --- |
|  |  | Bottom 80% | Top 20% |
| Set 1 (n=838) | Incorrect | 235 | 9 |
|  | Correct | 435 | 159 |
| Set 2 (n=778) | Incorrect | 177 | 4 |
|  | Correct | 445 | 152 |
| Set 3 (n=719) | Incorrect | 172 | 2 |
|  | Correct | 403 | 142 |
| Set 4 (n=526) | Incorrect | 129 | 3 |
|  | Correct | 291 | 103 |
| Set 5 (n=1,129) | Incorrect | 291 | 6 |
|  | Correct | 612 | 220 |

**Supplementary Table S2:**
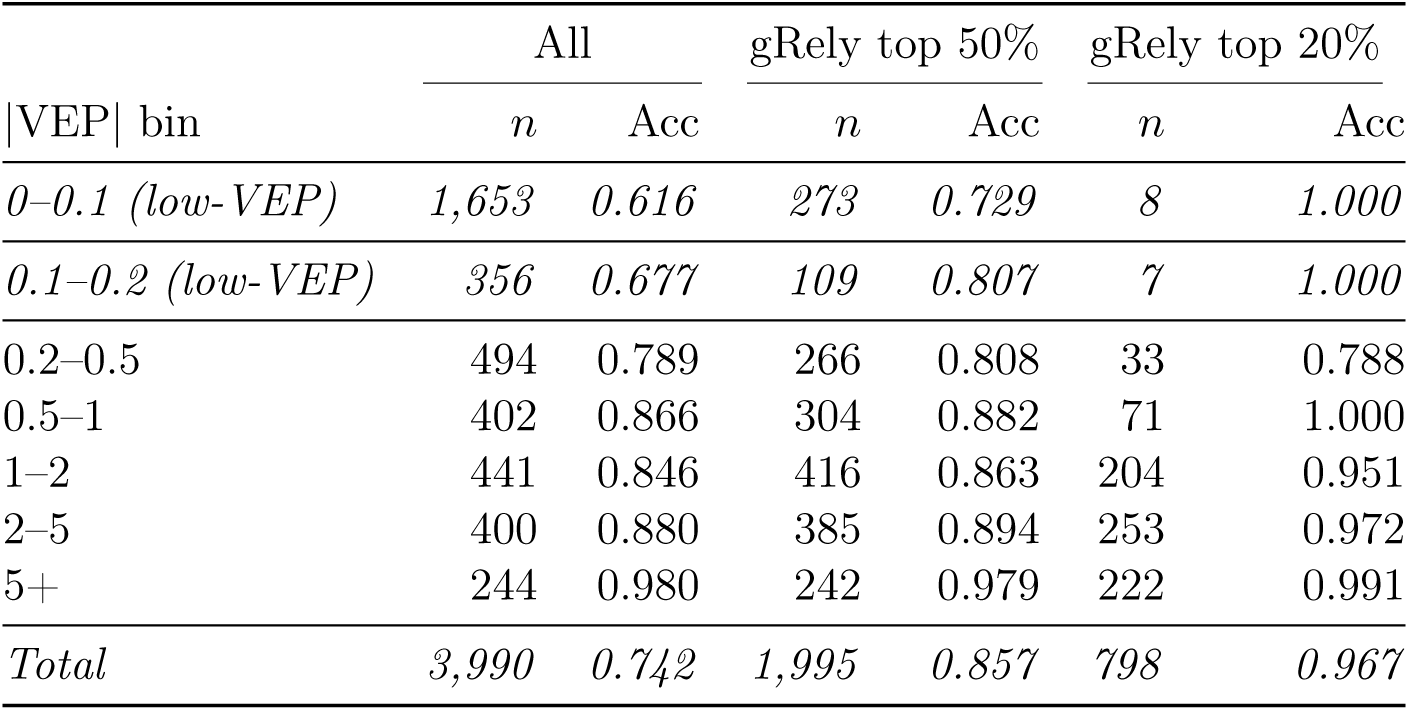
Sign concordance accuracy by VEP magnitude bin (test set, *n* = 3,990). For each |avg_logSum| bin, the number of test-set predictions and fraction with correct sign concordance are shown for all predictions, and after retaining the top 50% and top 20% of predictions by gRely score.

**Supplementary Table S3:** Mapping of GTEx tissues to Borzoi output tracks. For each tissue, the indices of the matching Borzoi GTEx RNA-seq tracks among the model’s 7,611 human output tracks. Predictions were averaged across tracks before scoring.

| Tissue | Borzoi tracks | Tissue | Borzoi tracks |
| --- | --- | --- | --- |
| adipose_subcutaneous | 7522, 7524 | liver | 7563, 7564, 7565 |
| adipose_visceral | 7523 | lung | 7566, 7567, 7568 |
| adrenal_gland | 7525, 7526, 7527 | minor_salivary_gland | 7586, 7587, 7588 |
| artery_aorta | 7535 | muscle | 7569, 7570, 7571 |
| artery_tibial | 7534, 7536 | nerve_tibial | 7572, 7573 |
| blood | 7532, 7533 | ovary | 7574, 7575, 7576 |
| brain_amygdala | 7540 | pancreas | 7577, 7578, 7579 |
| brain_cerebellar_hemisphere | 7539 | pituitary | 7580, 7581, 7582 |
| brain_cortex | 7541 | prostate | 7583, 7584, 7585 |
| breast | 7542, 7543, 7544 | skin_not_sun_exposed | 7589 |
| colon_sigmoid | 7548, 7549 | skin_sun_exposed | 7591 |
| colon_transverse | 7550 | small_intestine | 7592, 7593, 7594 |
| esophagus_gej | 7551, 7552 | spleen | 7595, 7596, 7597 |
| esophagus_mucosa | 7553 | stomach | 7598, 7599, 7600 |
| fibroblast | 7590 | testis | 7601, 7602 |
| heart_atrial_appendage | 7557 | thyroid | 7603, 7604, 7605 |
| heart_left_ventricle | 7558, 7559 | uterus | 7606, 7607 |
| kidney_cortex | 7560, 7561, 7562 | vagina | 7608, 7609, 7610 |
| LCL | 7531 |  |  |

**Supplementary Table S4:** Mapping of GTEx tissues to AlphaGenome output tracks. For each tissue, the index of the matching AlphaGenome `RNA_SEQ` output track.

| Tissue | Track | Tissue | Track |
| --- | --- | --- | --- |
| adipose_subcutaneous | 632 | liver | 600 |
| adipose_visceral | 658 | lung | 655 |
| adrenal_gland | 637 | minor_salivary_gland | 648 |
| artery_aorta | 611 | muscle | 659 |
| artery_tibial | 653 | nerve_tibial | 609 |
| blood | 661 | ovary | 596 |
| brain_amygdala | 618 | pancreas | 601 |
| brain_cerebellar_hemisphere | 634 | pituitary | 588 |
| brain_cortex | 615 | prostate | 636 |
| breast | 654 | skin_not_sun_exposed | 666 |
| colon_sigmoid | 603 | skin_sun_exposed | 643 |
| colon_transverse | 602 | small_intestine | 604 |
| esophagus_gej | 645 | spleen | 625 |
| esophagus_mucosa | 652 | stomach | 593 |
| fibroblast | 570 | testis | 591 |
| heart_atrial_appendage | 651 | thyroid | 624 |
| heart_left_ventricle | 650 | uterus | 598 |
| kidney_cortex | 606 | vagina | 599 |
| LCL | 567 |  |  |

**Supplementary Table S5:** Mapping of GTEx tissues to Enformer output tracks. Enformer has no GTEx RNA-seq outputs, so each tissue was mapped to a CAGE track. Three pairs of tissues share a track (colon_sigmoid and colon_transverse; esophagus_gej and esophagus_mucosa; skin_not_sun_exposed and skin_sun_exposed) and therefore receive identical VEPs for the same variant–gene pair.

| Tissue | Track | Tissue | Track |
| --- | --- | --- | --- |
| adipose_subcutaneous | 4902 | liver | 4686 |
| adipose_visceral | 4795 | lung | 4687 |
| adrenal_gland | 4977 | minor_salivary_gland | 4911 |
| artery_aorta | 4949 | muscle | 4691 |
| artery_tibial | 5164 | nerve_tibial | 5078 |
| blood | 4950 | ovary | 4688 |
| brain_amygdala | 5095 | pancreas | 4946 |
| brain_cerebellar_hemisphere | 4979 | pituitary | 5093 |
| brain_cortex | 5103 | prostate | 4690 |
| breast | 4976 | skin_not_sun_exposed | 4970 |
| colon_sigmoid | 4682 | skin_sun_exposed | 4970 |
| colon_transverse | 4682 | small_intestine | 4692 |
| esophagus_gej | 4683 | spleen | 4693 |
| esophagus_mucosa | 4683 | stomach | 4959 |
| fibroblast | 4908 | testis | 4694 |
| heart_atrial_appendage | 4975 | thyroid | 4696 |
| heart_left_ventricle | 4974 | uterus | 4910 |
| kidney_cortex | 4685 | vagina | 5175 |
| LCL | 5110 |  |  |

**Supplementary Figure S1:**
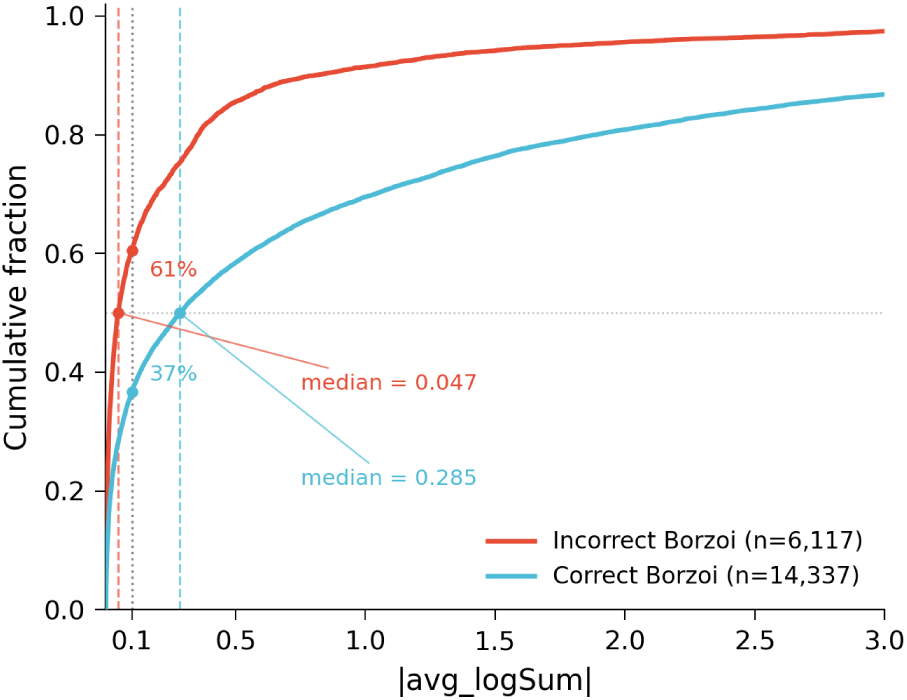
Predicted effect magnitude separates correct from incorrect Borzoi predictions only partially. Cumulative fraction (y-axis) of Borzoi predictions below a given predicted effect magnitude (|avg logSUM|; x-axis, truncated at 3) for sign-concordant (blue; *n* = 14,337) and sign-discordant (red; *n* = 6,117) predictions. Computed on the full labelled set (*n* = 20,454), not the pooled held-out test set used elsewhere.

**Supplementary Figure S2:**
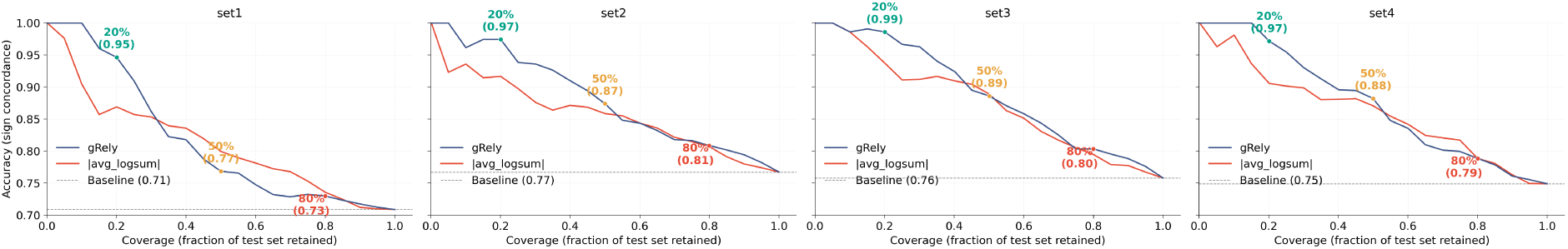
gRely exceeds magnitude filtering at low coverage in every held-out tissue-set fold. Sign concordance accuracy (y-axis) against coverage, the fraction of the test set retained (x-axis), for held-out predictions ranked by gRely score (blue) and by |avg logSUM| (red), for tissue-set folds set1–set4 (one panel each; set5 is shown in Fig 1). Dashed line, unfiltered per-fold baseline.

**Supplementary Figure S3:**
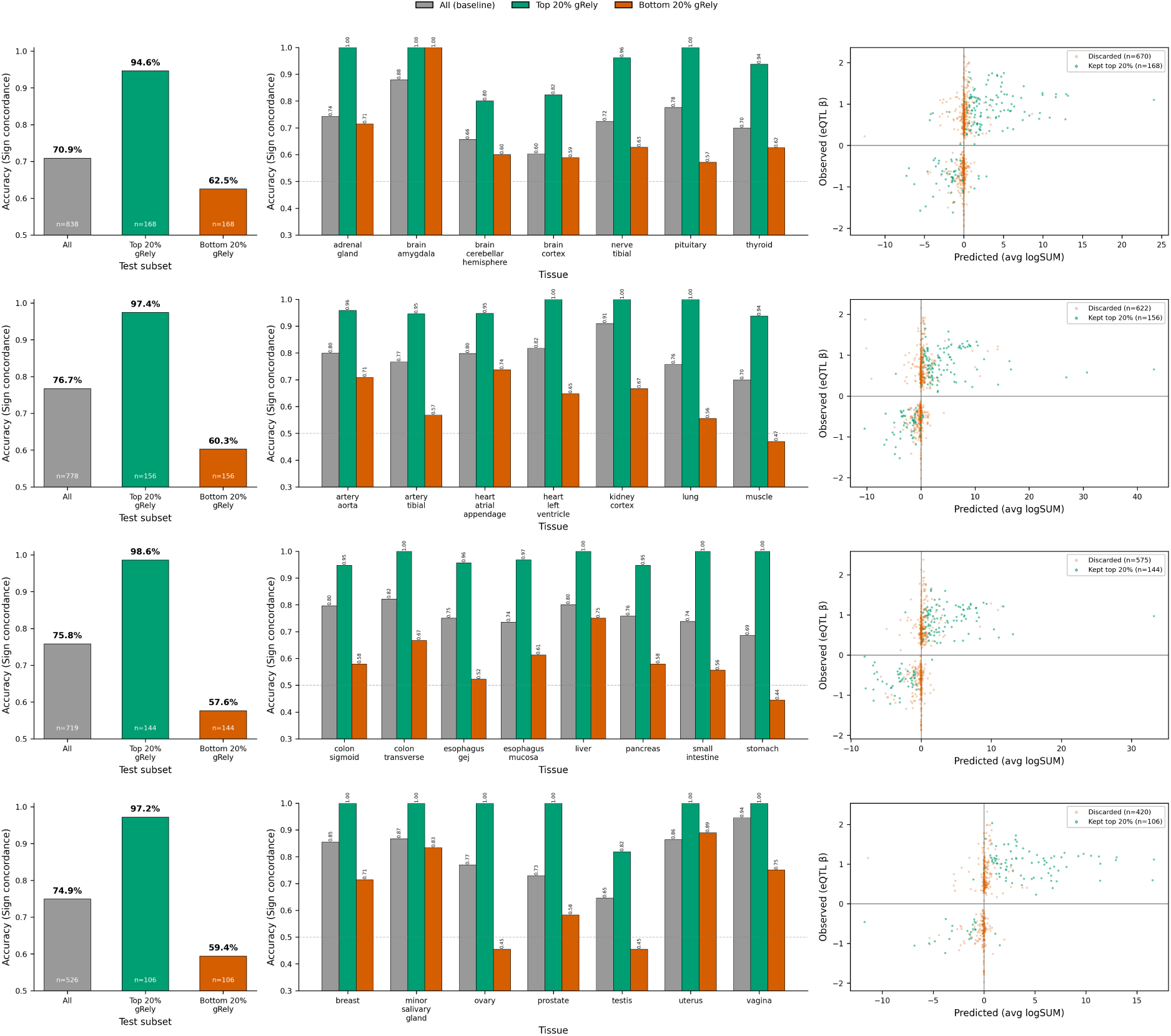
gRely separates reliable from unreliable Borzoi predictions in every held-out tissue-set fold. gRely applied to Borzoi VEPs on tissue-set folds set1–set4 (one fold per row; set5 is shown in Fig 1). Left: sign concordance (y-axis) for all predictions in the fold, the top 20% and the bottom 20% by gRely score (x-axis). Centre: the same three subsets (all, grey; top 20%, green; bottom 20%, orange) within each held-out tissue of that fold (x-axis). Right: observed eQTL effect size (*β*; y-axis) against predicted VEP (avg logSUM; x-axis) for predictions retained in the top 20% by gRely (green) and discarded (orange).

**Supplementary Figure S4:**
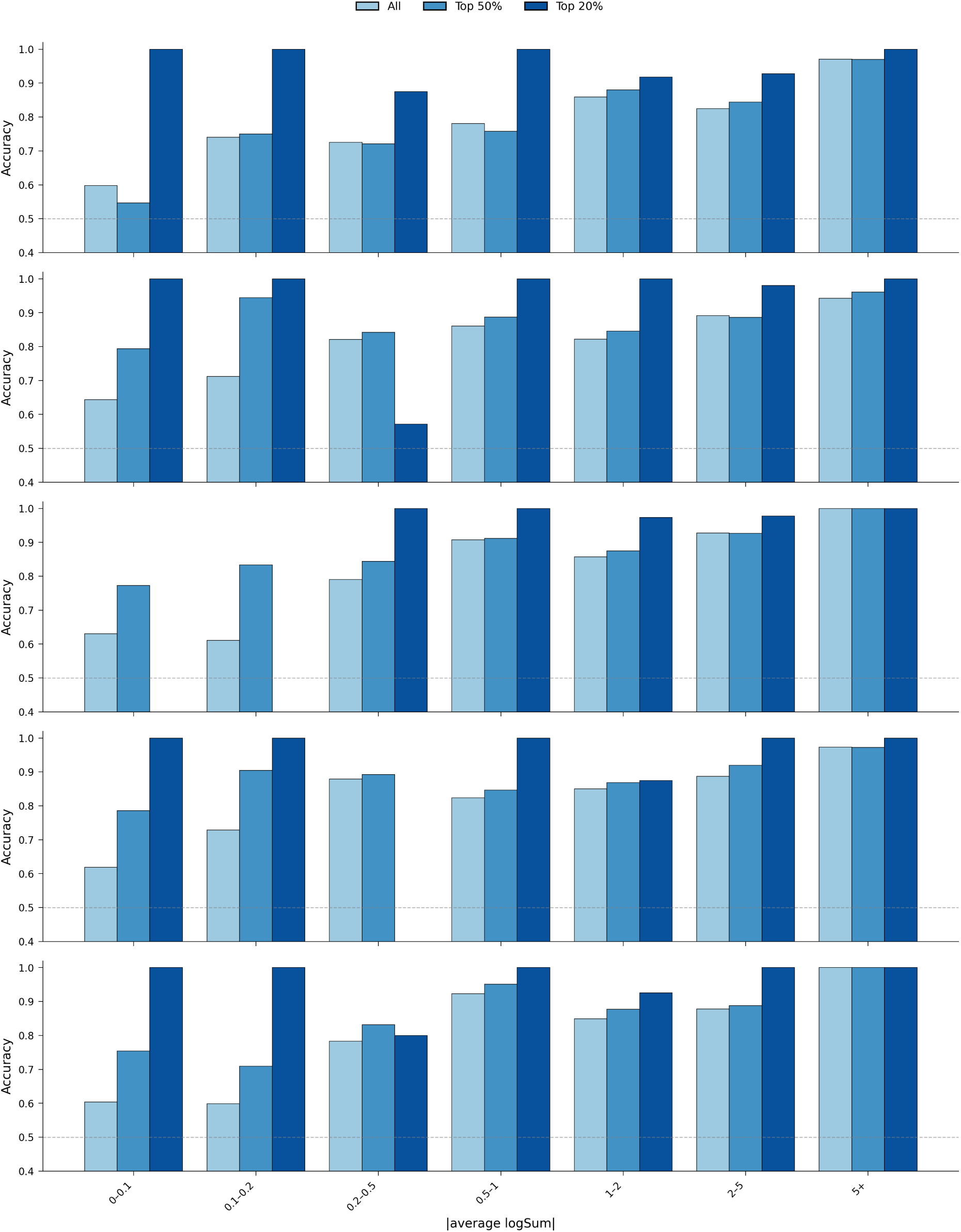
gRely raises accuracy within magnitude bins in every fold, with the largest gains in the low-VEP bins that magnitude thresholding discards. Sign concordance accuracy (y-axis) for held-out predictions binned by predicted effect magnitude (|avg logSUM|; x-axis) for all predictions in the bin (light blue), the top 50% by gRely score (medium blue) and the top 20% (dark blue); one row per held-out tissue-set fold (set1–set5, top to bottom).

**Supplementary Figure S5:**
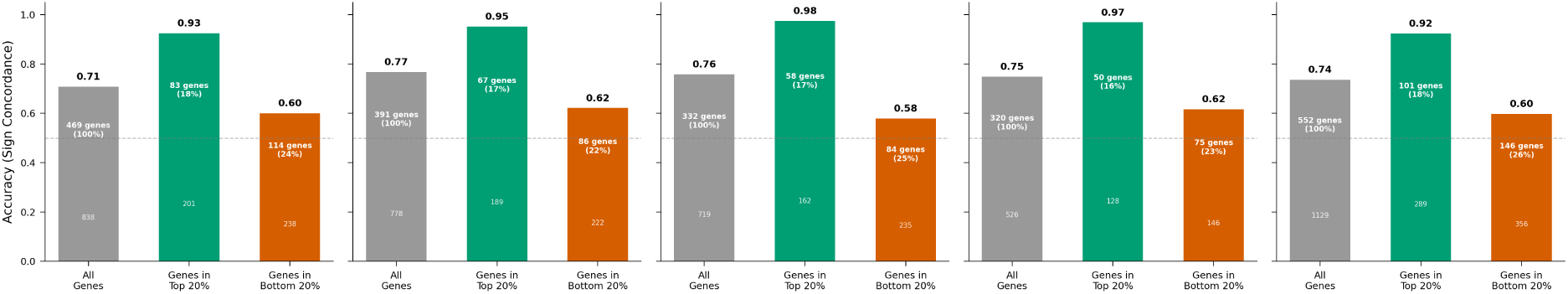
Borzoi is reliable for some genes and unreliable for others, and gRely separates the two. Sign concordance (y-axis) computed over *every* held-out prediction assigned to a gene, for all genes in the fold (grey), genes containing at least one prediction in the top 20% by gRely score (green), and genes containing at least one prediction in the bottom 20% (orange); one panel per held-out tissue-set fold (set1–set5). Genes carrying top-20% predictions are 16–18% of genes in a fold and reach 0.92–0.98 sign concordance, against per-fold baselines of 0.71–0.77; genes carrying bottom-20% predictions are 22–26% of genes and reach 0.58–0.62.

**Supplementary Figure S6:**
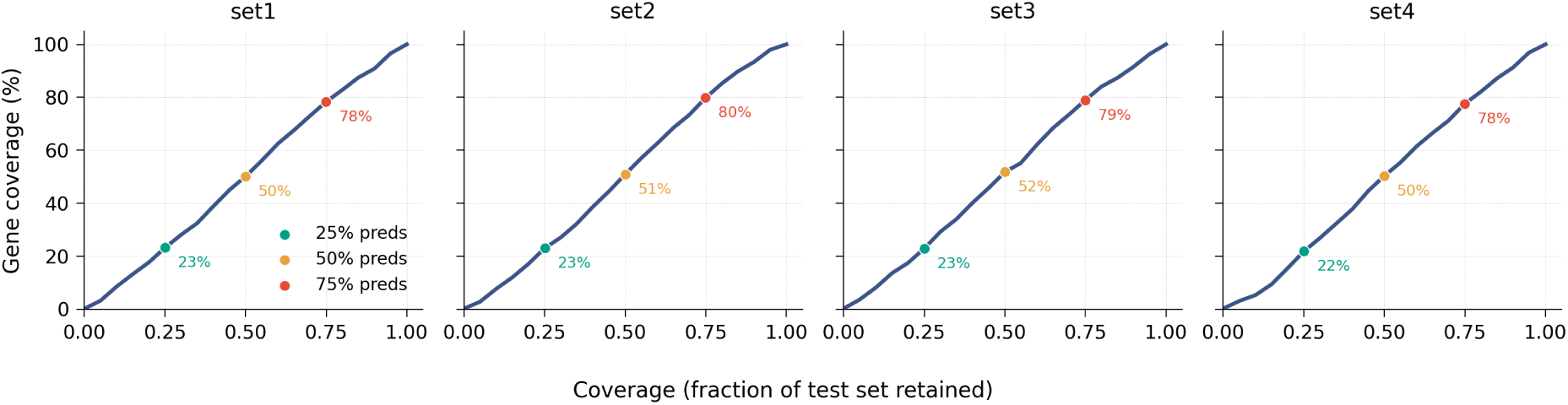
Filtering by gRely score retains genes in proportion to predictions rather than only retaining a few well-predicted loci. Percentage of unique genes retained (y-axis) at each prediction-level coverage threshold (x-axis), for held-out tissue-set folds set1–set4 (one panel each; set5 is shown in Fig 1).

**Supplementary Figure S7:**
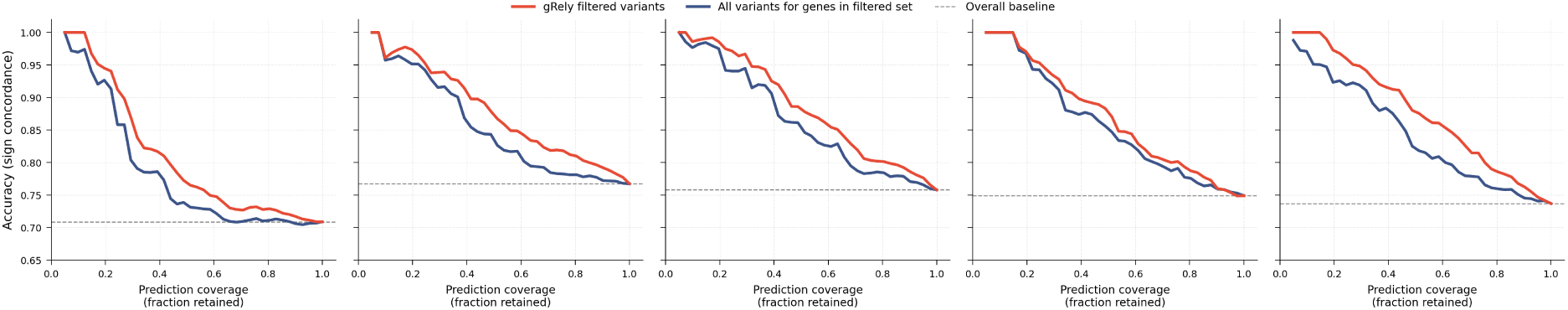
gRely can rank variants within a gene and not only genes against one another. Sign concordance (y-axis) against prediction coverage (x-axis) for the predictions gRely retains at each threshold (red) and for all predictions belonging to the genes those retained predictions come from (blue); one panel per held-out tissue-set fold (set1–set5). Dashed line, unfiltered per-fold baseline. Adding the discarded predictions of a retained gene back in lowers sign concordance at every coverage below 100%, in every fold.

**Supplementary Figure S8:**
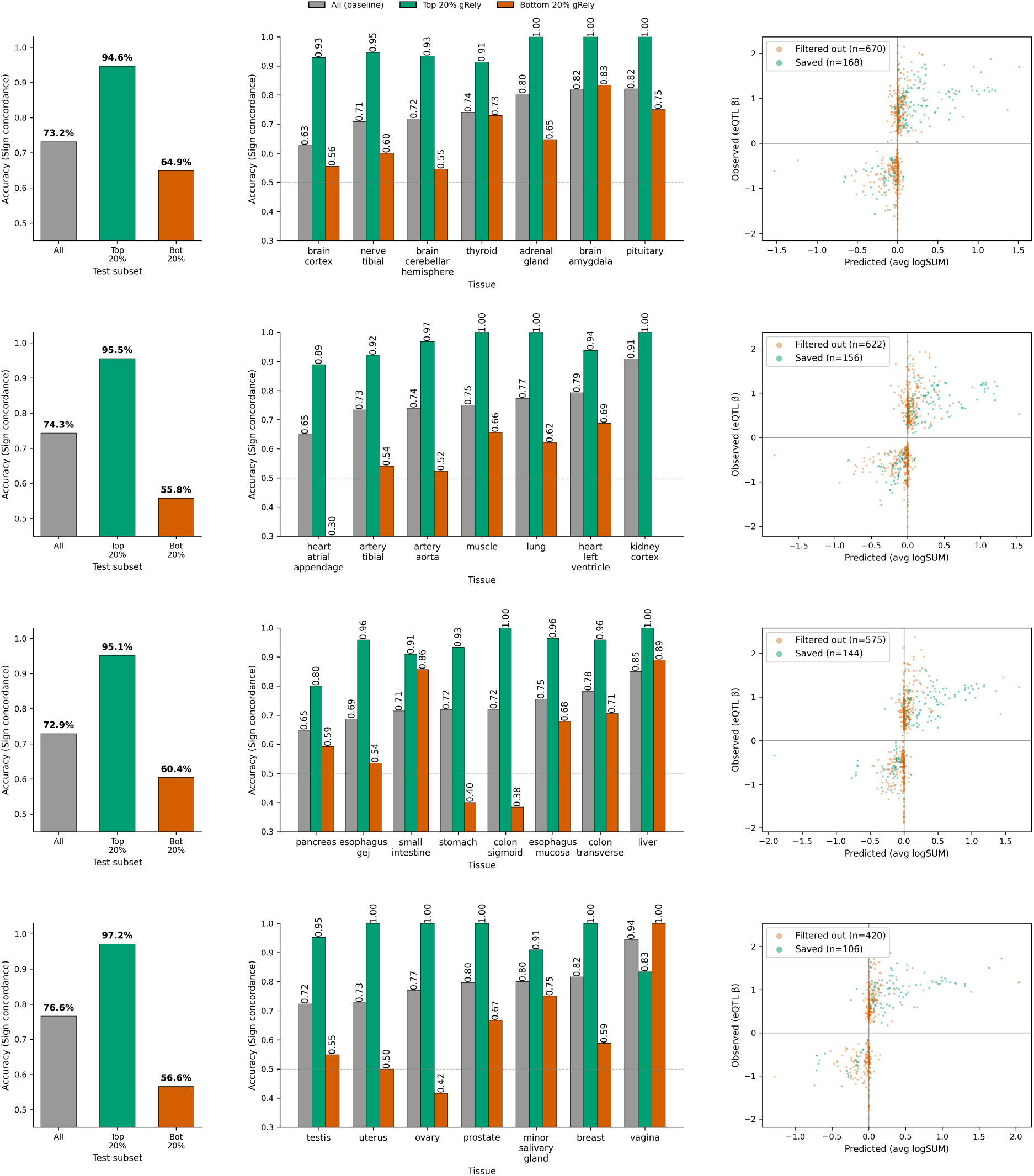
Reliability scores computed from Borzoi features transfer to AlphaGenome. gRely, trained on Borzoi VEPs and applied to AlphaGenome VEPs without retraining, evaluated on tissue-set folds set1–set4 (one fold per row; set5 is shown in main Fig 1). Left: sign concordance (y-axis) for all predictions in the fold, the top 20% and the bottom 20% by gRely score (x-axis). Centre: the same three subsets (all, grey; top 20%, green; bottom 20%, orange) within each held-out tissue (x-axis). Right: observed eQTL effect size (*β*; y-axis) against predicted VEP (avg logSUM; x-axis) for predictions retained in the top 20% (green) and discarded (orange).

**Supplementary Figure S9:**
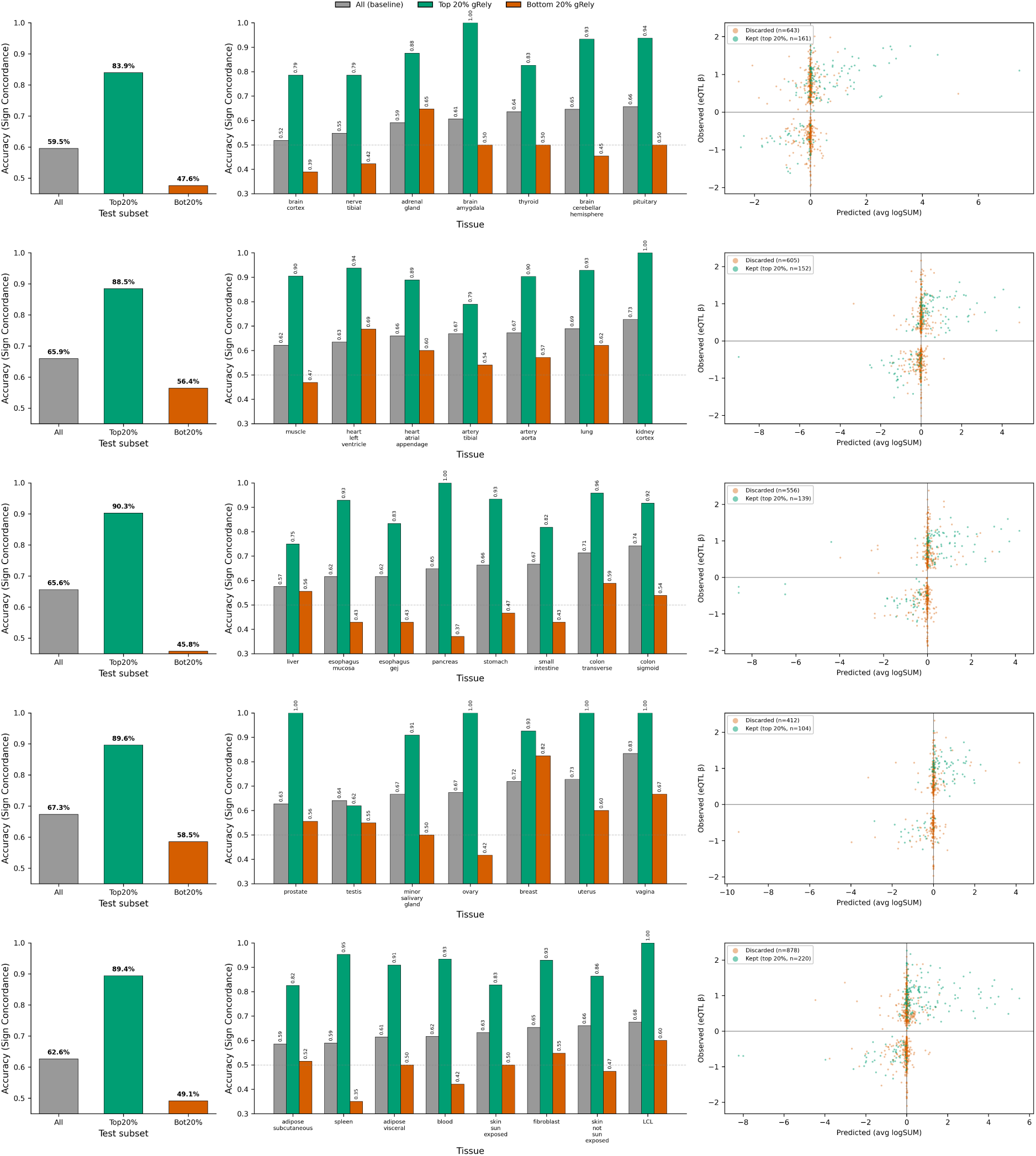
Reliability scores computed from Borzoi features transfer to Enformer, even though Enformer’s performance is well below Borzoi’s. gRely, trained on Borzoi VEPs and applied to Enformer VEPs without retraining, evaluated on all five held-out tissue-set folds (one fold per row, set1–set5). Left: sign concordance (y-axis) for all predictions in the fold, the top 20% and the bottom 20% by gRely score (x-axis). Centre: the same three subsets (all, grey; top 20%, green; bottom 20%, orange) within each held-out tissue (x-axis). Right: observed eQTL effect size (*β*; y-axis) against predicted VEP (avg logSUM; x-axis) for the top 20% (green) and the discarded predictions (orange).

**Supplementary Figure S10:**
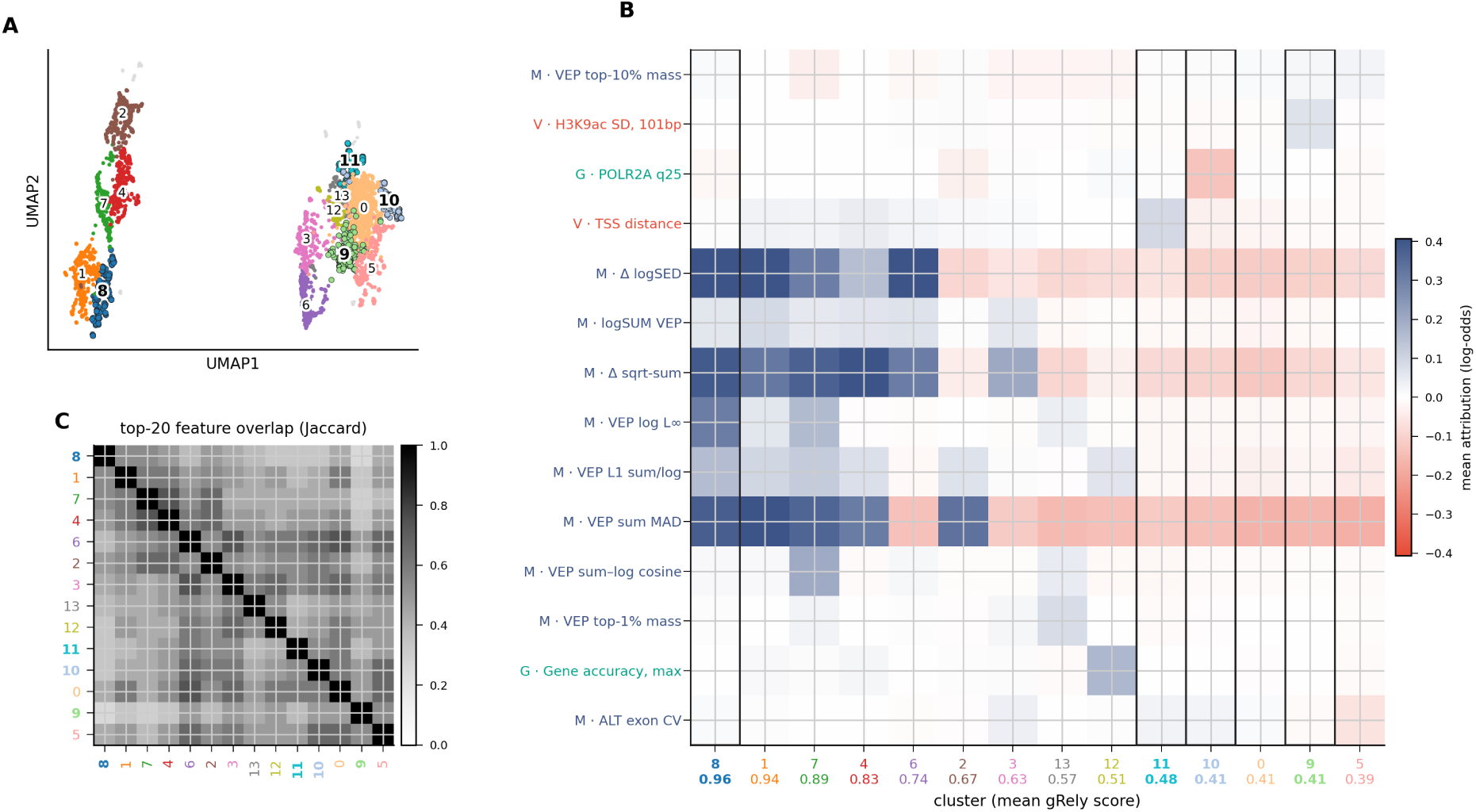
Clustering attribution scores identify distinct set of features driving reliability score. Leiden clustering of the SHAP attribution embedding on the pooled held-out test set (*n* = 3,990 predictions × 1,161 features). **A.** UMAP embedding of per-prediction attributions, coloured by Leiden cluster. **B.** Mean SHAP attribution (log-odds; colour scale, blue positive, red negative) for the highest-ranked features (y-axis) in each cluster (x-axis), with clusters ordered by mean gRely score (annotated below each column, 0.96 to 0.39). Feature labels are prefixed by domain (M, model; V, variant; G, gene). **C.** Jaccard overlap between the top-20 attributed features of each pair of clusters (colour scale). Clusters share features partially and no two clusters have the same top-20 set.

**Supplementary Figure S11:**
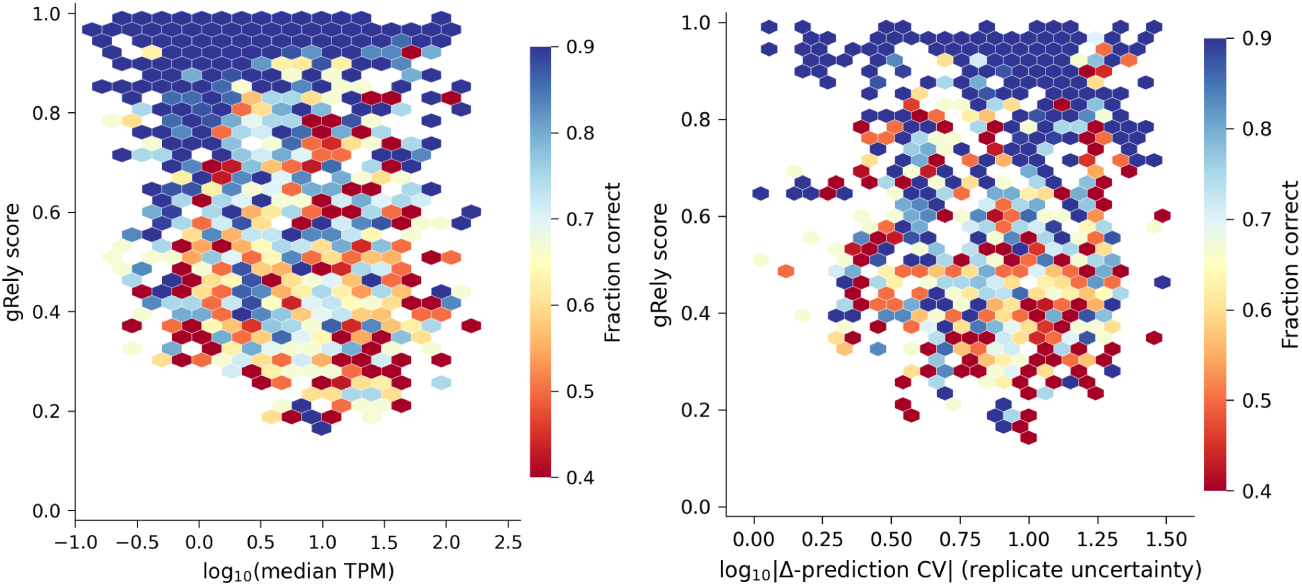
Features ranked highly by gRely do not separate correct from incorrect predictions on their own. Hexagonal bins of held-out predictions (pooled test set, *n* = 3,990) with gRely score (y-axis) against a single feature (x-axis), coloured by the fraction of predictions in each bin with sign-concordant Borzoi VEP (colour scale). Left: observed median TPM of the target gene in the target tissue (log_10_). Right: replicate uncertainty, the coefficient of variation of the (alt − ref) prediction across Borzoi replicates (log_10_). Each feature tracks the gRely score loosely (Spearman *ρ* = −0.17 and +0.13 respectively), and the fraction correct varies along the score axis rather than along the feature axis.

**Supplementary Figure S12:**
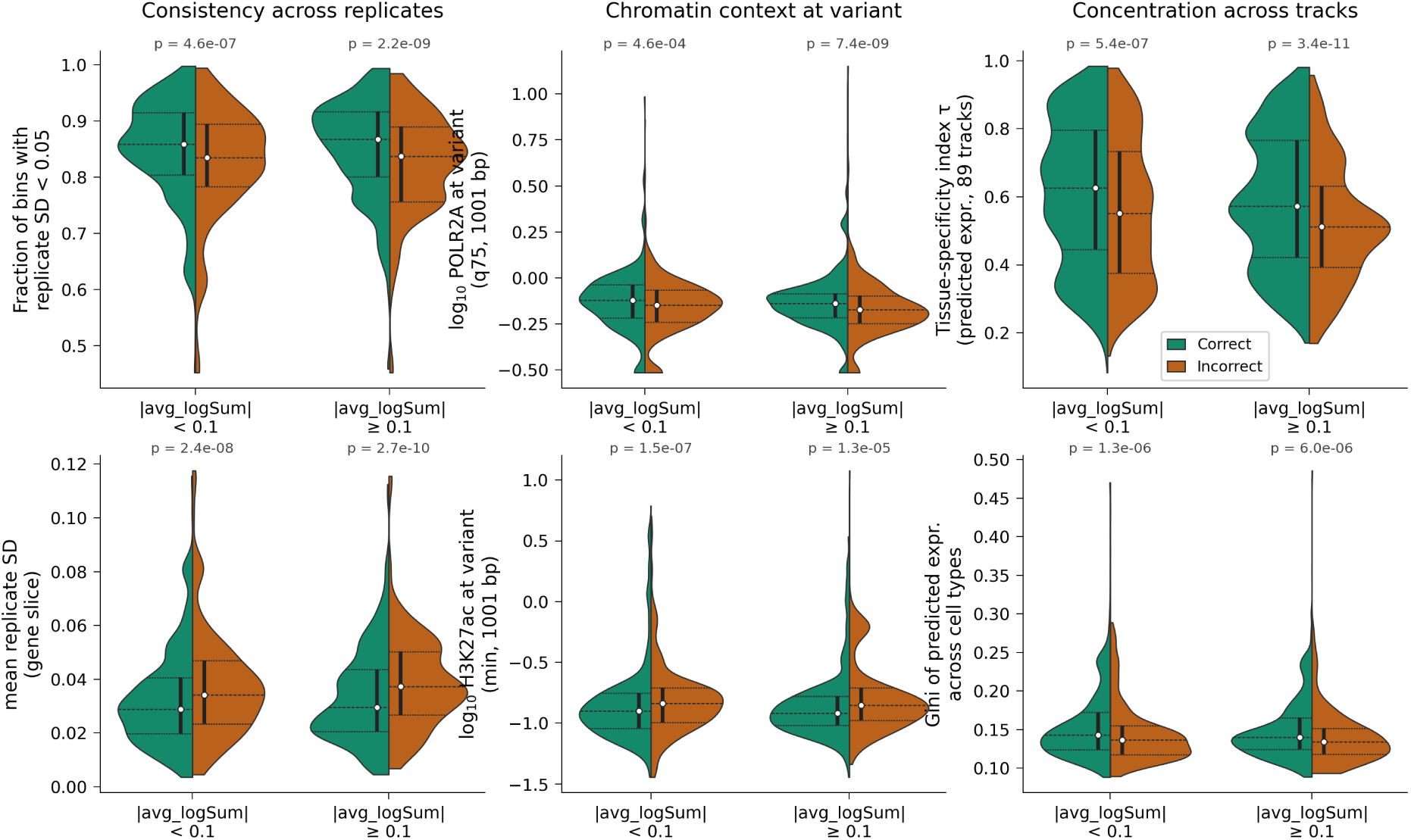
Replicate consistency, chromatin at the variant, and concentration of the predicted signal separate correct from incorrect Borzoi predictions within both magnitude strata. Distributions of six features for sign-concordant (green) and sign-discordant (orange) Borzoi predictions, on the pooled held-out test set (*n* = 3,990; *n* = 1,653 in the low-VEP stratum). Each panel shows one feature (y-axis) stratified by predicted effect magnitude (|avg logSUM| < 0.1 and ≥ 0.1; x-axis), with two features per signal class (columns). Consistency across replicates (left column): fraction of gene-body bins with cross-replicate standard deviation < 0.05 (top) and mean cross-replicate standard deviation over the gene slice (bottom). Chromatin context at the variant (centre column): log_10_ POLR2A signal (75th percentile, 1,001 bp window; top) and log_10_ H3K27ac signal (minimum, 1,001 bp window; bottom). Concentration across tracks (right column): tissue-specificity index *τ* of the predicted expression across the 89 GTEx tracks, over exonic bins (top), and Gini coefficient of the predicted expression across cell types (bottom). Correct predictions show higher cross-replicate agreement and more concentrated predicted signal in both strata, so the separation is not effect magnitude in another form. The distributions overlap throughout, which is why gRely aggregates these features rather than thresholding on any one of them (Supplementary Fig. S11).

**Supplementary Figure S13:**
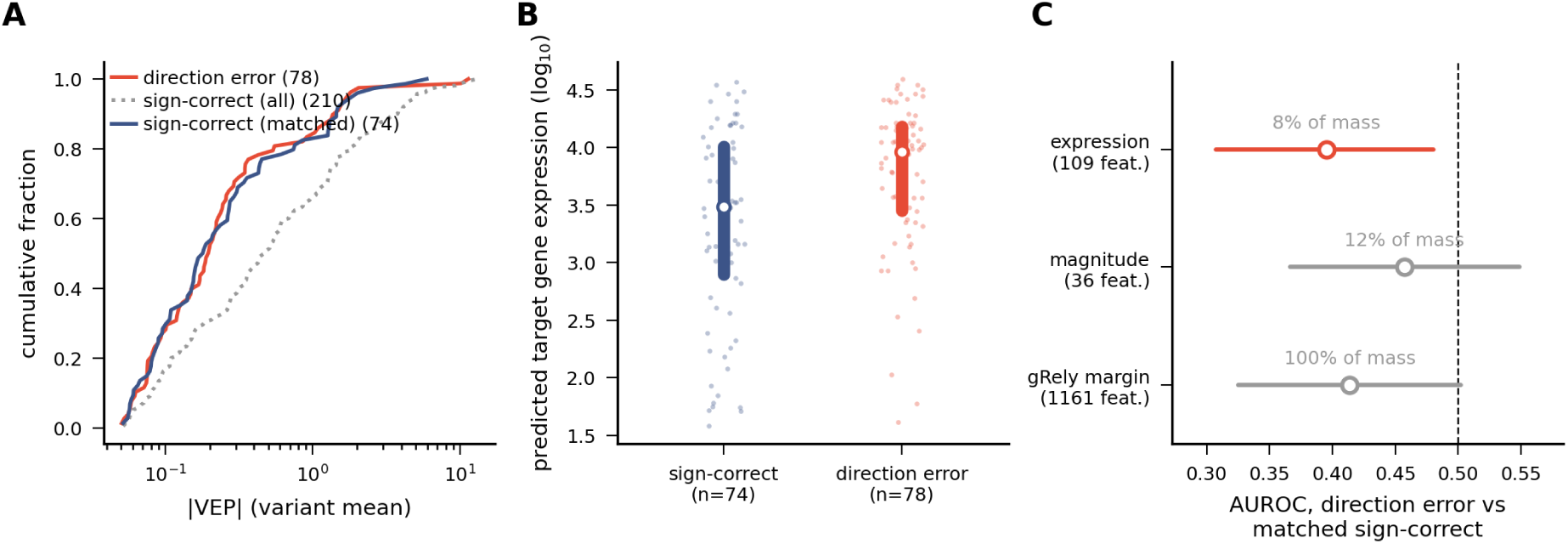
Direction errors occur at more highly expressed target genes. Direction errors are held-out predictions for which Borzoi ranks the effect magnitude correctly but predicts the wrong sign. Direction errors (*n* = 78) are compared with sign-correct predictions matched on |VEP| at the variant level (*n* = 74), so the two groups differ in direction alone. **A.** Cumulative fraction (y-axis) against |VEP| (variant mean; x-axis, log scale) for direction errors (red), all sign-correct predictions (dotted) and the matched sign-correct set (blue). The matched set overlays the direction errors, confirming that matching removes the magnitude difference. **B.** Predicted target gene expression (log_10_; y-axis) for matched sign-correct predictions and direction errors (x-axis). **C.** AUROC (x-axis) separating direction errors from matched sign-correct predictions, where the score being ranked is the sum of *signed* SHAP values over a family of gRely features: expression-family (109 features), magnitude-family (36), or all features (1,161), the last being equivalent to gRely’s margin and therefore to its reliability score. The orientation is inverted relative to **A** and **B**. Values below the dashed line at 0.5 mean the family contributes *more negatively* in direction errors, that is, it lowers gRely’s reliability score for them; a value of 0.5 means no difference. Expression-family features are shifted downward in direction errors, consistent with **B**, and account for essentially all of the shift seen in the full margin despite carrying [8]% of attribution mass. The magnitude family is matched away by construction in **A**, so its flat result is expected.

**Supplementary Figure S14:**
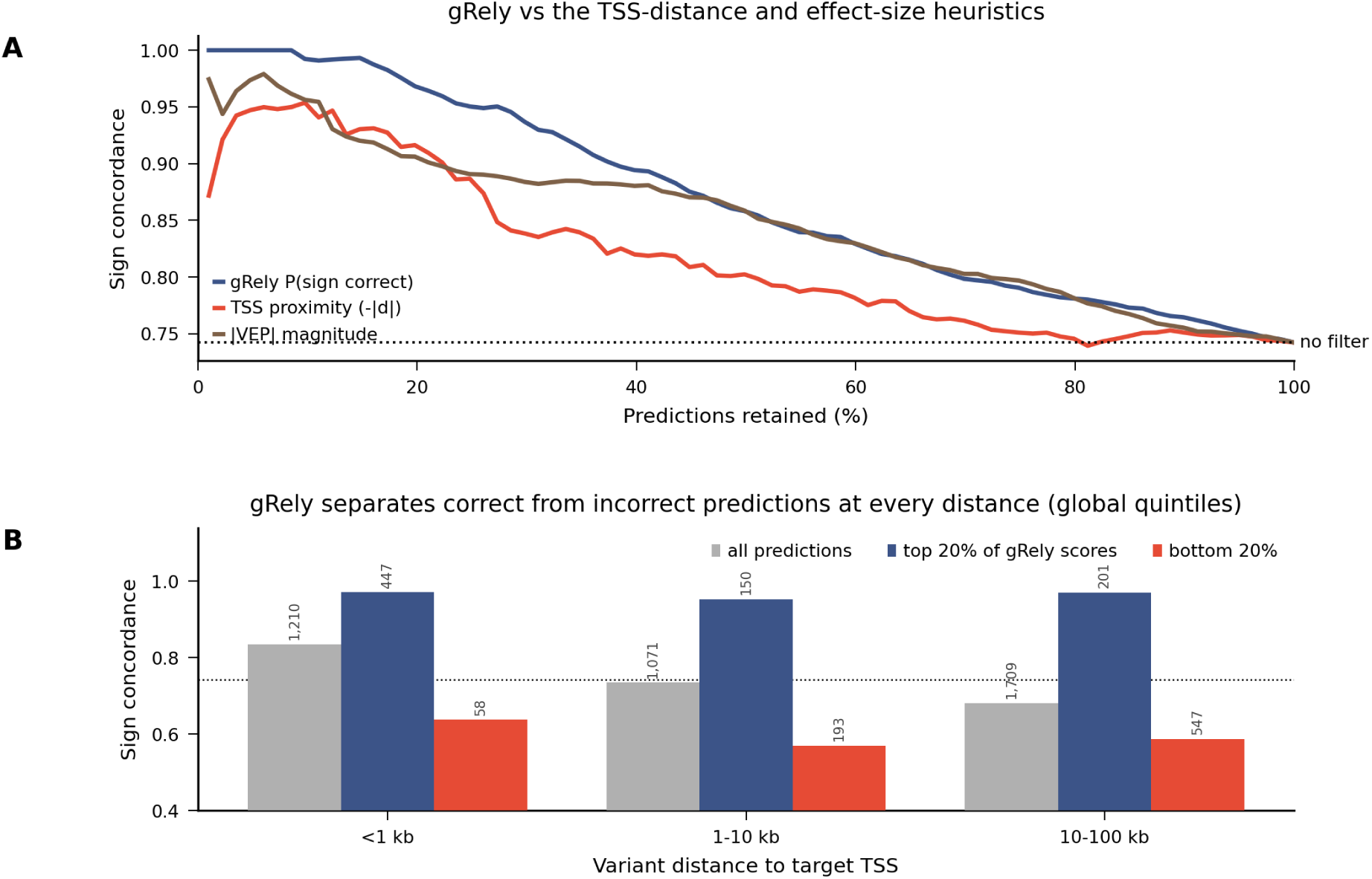
gRely can score predictions at varying distance from the TSS. **A**, sign concordance among retained predictions (y-axis) as a function of the fraction of held-out predictions retained (x-axis), ranking predictions by gRely reliability score (blue), by proximity to the target TSS (red), or by |VEP| magnitude (brown); the dotted line is concordance with no filtering. gRely retains more concordant predictions than either heuristic. **B**, sign concordance by variant distance to the target TSS, for all held-out predictions in the bin (grey), for those in the top quintile of gRely scores (blue), and for those in the bottom quintile (red); quintile cuts are global over the held-out set rather than within bins, so the number of predictions surviving each cut varies by bin (*n*, annotated above each bar). The dotted line is overall concordance with no filtering. Baseline concordance declines with distance, reproducing the known distal failure mode, but the gRely quintiles separate correct from incorrect predictions within every stratum. Variant–gene pairs are restricted to genes with exons inside Borzoi’s central prediction window, so no pair exceeds *∼*98 kb and the most distal regime cannot be assessed here.

**Supplementary Figure S15:**
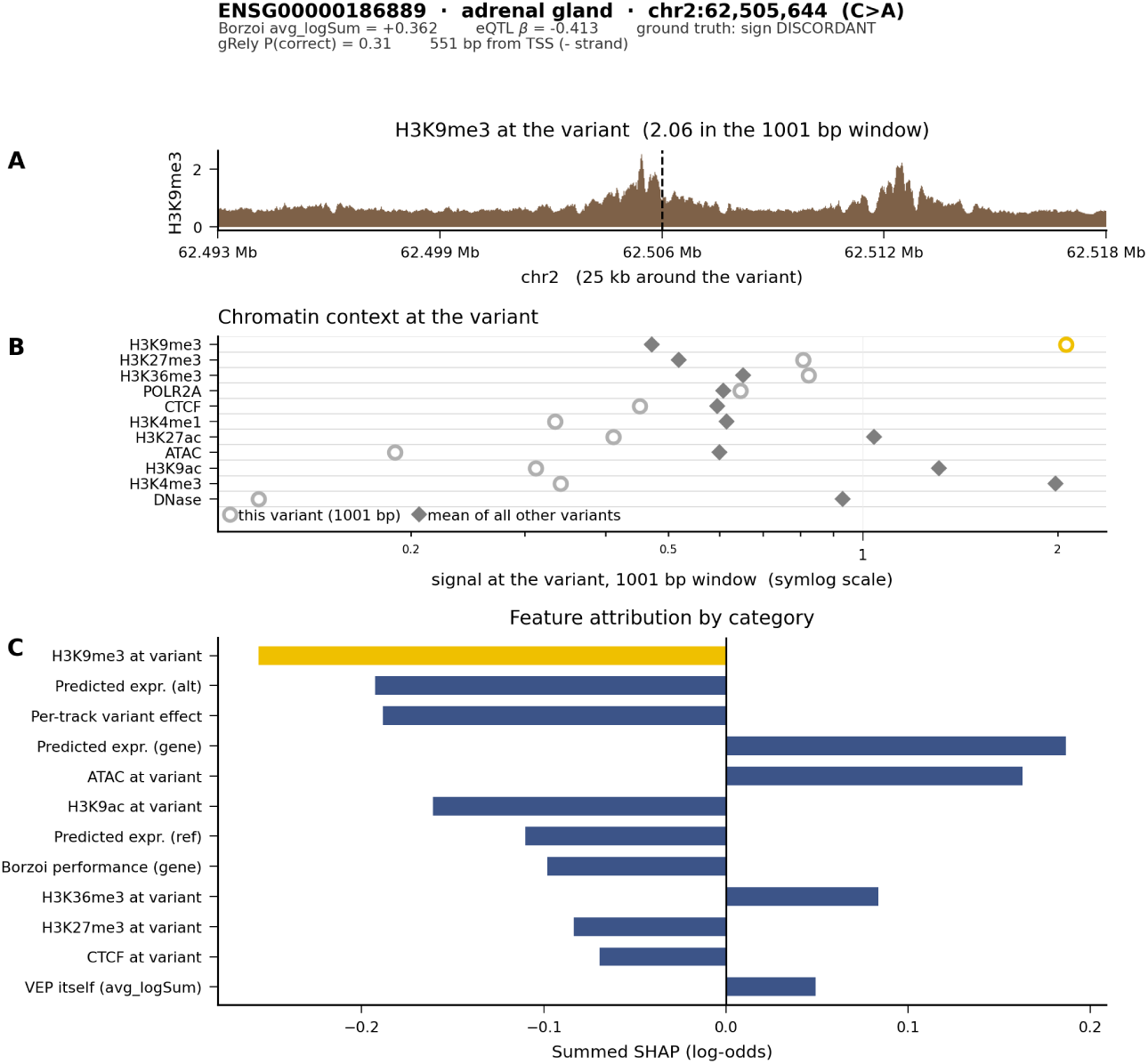
For every prediction, gRely returns a reliability score together with the evidence that produced it. Readout for a single (variant, gene, tissue) prediction: variant chr2:62,505,644 (C>A), gene *ENSG00000186889*, adrenal gland. The prediction and its ground truth: Borzoi avg_logSum = +0.362 against a fine-mapped eQTL effect of *β* = −0.413, so the direction of effect is discordant despite the variant lying only 551 bp from the target TSS. gRely assigns this prediction *P* (correct) = 0.31, flagging it as unreliable. **A** H3K9me3 coverage (y-axis) over 25 kb centred on the variant (x-axis, genomic position; dashed line, variant position). **B** Mean signal in a 1001 bp window centred on the variant (x-axis, log) for eleven chromatin tracks, comparing this variant (open circles) with the mean over all other test variants (grey diamonds); H3K9me3 is highlighted (yellow). The variant sits in repressed, inaccessible chromatin: H3K9me3 is the highest of the eleven marks here, together with H3K27me3 and H3K36me3, while DNase, ATAC, H3K4me3, H3K9ac and H3K27ac all fall below it. **C** SHAP attribution for this prediction summed within feature category (x-axis, log-odds), top twelve categories by absolute contribution, with H3K9me3 at the variant highlighted (yellow). H3K9me3 at the variant carries the largest negative contribution, ahead of the predicted-expression and per-track variant-effect categories. The reduced score is therefore traceable to a named contextual feature, and the same readout is available for any (variant, gene, tissue) tuple gRely scores.

